# Spatial Mark-Resight for Categorically Marked Populations with an Application to Genetic Capture-Recapture

**DOI:** 10.1101/299982

**Authors:** Ben C. Augustine, Frances E. C. Stewart, J. Andrew Royle, Jason T. Fisher, Marcella J. Kelly

## Abstract

The estimation of animal population density is a fundamental goal in wildlife ecology and management, commonly met using mark recapture or spatial mark recapture (SCR) study designs and statistical methods. Mark-recapture methods require the identification of individuals; however, for many species and sampling methods, particularly noninvasive methods, no individuals or only a subset of individuals are individually identifiable. The unmarked SCR model, theoretically, can estimate the density of unmarked populations; however, it produces biased and imprecise density estimates in many sampling scenarios typically encountered. Spatial mark-resight (SMR) models extend the unmarked SCR model in three ways: 1) by introducing a subset of individuals that are marked and individually identifiable, 2) introducing the possibility of individual-linked telemetry data, and 3) introducing the possibility that the capture-recapture data from the survey used to deploy the marks can be used in a joint model, all improving the reliability of density estimates. The categorical spatial partial identity model (SPIM) improves the reliability of density estimates over unmarked SCR along another dimension, by adding categorical identity covariates that improve the probabilistic association of the latent identity samples. Here, we combine these two models into a “categorical SMR” model to exploit the benefits of both models simultaneously. We demonstrate using simulations that SMR alone can produce biased and imprecise density estimates with sparse data and/or when few individuals are marked. Then, using a fisher (*Pekania pennanti*) genetic capture-recapture data set, we show how categorical identity covariates, marked individuals, telemetry data, and jointly modeling the capture survey used to deploy marks with the resighting survey all combine to improve inference over the unmarked SCR model. As previously seen in an application of the categorical SPIM to a real-world data set, the fisher data set demonstrates that individual heterogeneity in detection function parameters, especially the spatial scale parameter *σ*, introduces positive bias into latent identity SCR models (e.g., unmarked SCR, SMR), but the categorical SMR model provides more tools to reduce this positive bias than SMR or the categorical SPIM alone. We introduce the possibility of detection functions that vary by identity category level, which will remove individual heterogeneity in detection function parameters than is explained by categorical covariates, such as individual sex. Finally, we provide efficient SMR algorithms that accommodate all SMR sample types, interspersed marking and sighting periods, and any number of identity covariates using the 2-dimensional individual by trap data in conjunction with precomputed constraint matrices, rather than the 3-dimensional individual by trap by occasion data used in SMR algorithms to date.

## 1 Introduction

The estimation of animal population density is a fundamental problem in wildlife ecology and management (Krebs, 1972) and increasingly, population density is estimated using noninvasive methods such as remote cameras (Burton *et al.*, 2015) and the collection of genetic material from scat or hair samples (Waits & Paetkau, 2005). These methods allow for systematic data collection over large areas with less effort than live capture surveys (Kelly *et al.*, 2012). One of the most robust and well-developed class of models for estimating density, mark-recapture, and its extension to spatial mark-recapture (SCR), require the determination of the individual identities of all captured individuals; however, many species are not reliably identifiable using noninvasive methods. SCR models specifically, have been extended to accommodate fully unmarked populations (Chandler & Royle, 2013), but unfortunately, the unmarked SCR model produces biased and imprecise density estimates in many scenarios typically encountered when using noninvasive methods (Chandler & Royle, 2013; Augustine *et al.*, 2018). The SCR modeling framework does; however, offer several ways to improve density estimates for unmarked populations including the introduction of marked individuals (Chandler & Royle, 2013; Sollmann *et al.*, 2013), informative priors related to home range size (Chandler & Royle, 2013; Ramsey *et al.*, 2015), telemetry data (Sollmann *et al.*, 2013; Whittington *et al.*, 2016), and/or partial identity information (e.g., categorical marks such as individual sex; Augustine *et al.*, 2018). SCR models utilizing only partial individual identities have only recently been introduced and have not yet been extended to accommodate marked individuals or individual-linked telemetry data. Combining all these data sources into a single model to capitalize on as much information about detected animals as possible to improve density estimates is the subject of this paper.

The main features of an SCR model are: 1) an explicit model for the distribution of individual activity centers across a state-space (i.e., the study area) and 2) a model for trap level detection probability that is a decreasing function of the distance between individual activity centers and the trap locations. The detection function typically has two parameters, the first of which is either λ_0_ or *p*_0_, the baseline parameters for the expected number of counts (detection rate) or the detection probability for a trap located exactly at an activity center, respectively, depending on if the observation model is Poisson or Bernoulli. The second detection function parameter is *σ* which determines how quickly detection rate or detection probability declines with the distance between an activity center and a trap, and is related to the home range size of the species under study (Royle *et al.*, 2013). When summed over the capture occasions, the observed data are the individual by trap number of counts or detections.

The unmarked SCR model is an SCR model where the individual identities are not observable, leading to trap-referenced, unidentified counts. The inferential process of unmarked SCR is to probabilistically resolve the true individual identities of the observed trap-referenced counts and then incorporate the uncertainty in individual identity into the estimates of abundance and density. The process by which the latent individual identities are probabilistically resolved involves relating the latent identity trap-referenced counts to the latent activity center structure of the SCR process model, and then updating the latent identities of these samples in a Markov chain Monte Carlo (MCMC) algorithm. This process uses the spatial location where samples were collected to probabilistically link samples collected closer together in space more frequently than those collected further apart, with the estimated *σ* parameter playing a large role in determining how frequently any two latent identity samples should be combined given their spatial proximity. Thus, the *σ* parameter estimate is particularly important for correctly allocating the latent identity samples to the latent activity centers (Chandler & Royle, 2013; Augustine *et al.*, 2018), and unmarked SCR estimates can be improved by using informative priors for *σ* and/or telemetry data (Chandler & Royle, 2013).

As classified by Augustine *et al.* (2018), unmarked SCR is the most basic type of spatial partial identity model (SPIM), where a SPIM is defined to be a partial identity model that uses the spatial location where samples were collected to improve the probabilistic identity associations among samples. A second important feature of a SPIM is that it capitalizes on two key ecological spatial concepts to resolve the uncertainty in individual identities – population density and home range size. Augustine *et al.* (2018) demonstrate that when sample identities are latent, the magnitude of uncertainty in individual identity is larger in higher density populations and/or when home ranges are larger, as quantified by the *σ* parameter in SCR models. Augustine *et al.* (2018) propose using a metric based on the Simpsons Diversity Index (Simpson, 1949) to quantify the expected magnitude of uncertainty in individual identity for a given density and *σ*, which they call the “Individual Diversity Index” (IDI). Further, they propose that the realized magnitude of uncertainty in individual identity in a SPIM can be quantified by the uncertainty in *n^cap^*, the number of individuals captured. This is a known quantity in a regular mark recapture model, but is a a random variable in a SPIM, and the uncertainty in *n^cap^* correlates well with the magnitude of uncertainty in abundance and density estimates (Augustine *et al.*, 2018).

Two other SPIMs, spatial mark-resight (SMR Sollmann *et al.*, 2013) and the categorical SPIM (Augustine *et al.*, 2018), have extended unmarked SCR to incorporate additional individually identifying information and improve density estimation with latent identity samples. First, spatial mark-resight introduces the possibility of deterministic identity associations between the marked and identifiable individual samples, and introduces more deterministic constraints for the latent identity samples if the marked status of some or all samples can be observed. Second, the categorical SPIM uses partially identifying categorical covariates to reduce the uncertainty in the individual identities of latent identity samples by adding deterministic constraints between samples whose categorical identities conflict, and by improving the probabilistic identity associations using the frequencies with which category levels are found in the population. We will discuss each of these SPIMs in more detail and then propose that they be combined into a “categorical SMR” model that uses all of these sources of information for improved density estimation with latent identity samples.

If a portion of a wildlife population is either naturally marked (e.g., unique markings or morphological features for some individuals, e.g. Rich *et al.*, 2014) or marks are deployed before the capture-recapture survey (e.g., VHF or GPS tracking collars, ear tags, etc. Sollmann *et al.*, 2013; Whittington *et al.*, 2016), spatial mark-resight models can exploit this information to improve density estimation over an unmarked SCR model. The SMR model allows for the marked portion of the population to be individually identifiable so that some or all of their samples can be deterministically linked together, reducing the number of samples whose individual identities need to be probabilistically resolved. The samples that cannot be deterministically linked are then treated like the latent identity samples in unmarked SCR that must be probabilistically associated using spatial information. The main advantage of SMR is the availability of deterministic linkages provided by the marked and identifiable samples, but the presence of marked individuals introduces identity constraints that can be used to better resolve the uncertainty in the latent identity samples. In SMR, there are three classes of latent identity sample types carrying different amounts of individually-identifying information (as described in Royle *et al.*, 2013). The least informative type of sample is one for which the marked status cannot be determined. These samples must be probabilistically associated with both marked and unmarked individuals and are equivalent to a latent identity sample in unmarked SCR. Marked but not individually identifiable and unmarked samples are more informative because they can only be probabilistically associated with the marked and unmarked individuals, respectively, reducing the number of possible individuals a latent identity sample could belong to. A second advantage of SMR is that telemetry can be linked to specific marked individuals which improves the estimates of three parameters: 1) the activity center locations of the telemetered individuals (e.g., Whittington *et al.*, 2016), 2) the parameters of more ecologically-informed detection function models (e.g., Linden *et al.*, 2017), and 3) *σ* which is the only parameter that may be informed by telemetry in unmarked SCR.

Recently introduced generalized SMR (Whittington *et al.*, 2016), distinguished from conventional SMR (Sollmann *et al.*, 2013) described above, provides an additional source of information that improves the probabilistic association of latent identity samples. The main purpose of generalized SMR is to correctly describe the distribution of marked individual activity centers across the state-space. An important assumption in conventional SMR is that the spatial distribution of activity centers for the marked individuals is either the same as the spatial distribution of the unmarked individuals (e.g., the marked individuals constitute a random sample from the state-space) or that the spatial distribution of marked individuals can be specified using a parametric model (Royle *et al.*, 2013). However, generally, the information necessary for specifying the spatial distribution for the marked individuals is either not available or imprecise. Generalized SMR determines the spatial distribution of the marked individuals by modeling the process by which animals came to be marked (the “marking process”), improving inference over models where the spatial distribution of the marked individuals is not correctly specified. A second advantage of modeling the marking process is that because all captured individuals are individually identifiable during marking, additional information about the detection function parameters and the locations of the marked and unmarked individual activity centers is provided, allowing for the identities of the latent identity samples to be probabilistically resolved with less uncertainty and for density to be estimated with more accuracy and precision.

Augustine *et al.* (2018) introduced the categorical SPIM where, rather than individuals being fully unmarked, they are categorically marked, precluding the determination of individual identification, but introducing some information to reduce the uncertainty in the probabilistic identity associations among samples and thus producing more reliable and precise density estimates. To illustrate this model, consider that identity covariates can be obtained from noninvasive samples; for example, sex, age class, color morph, and/or antler features may be determined from remote camera surveys or locus-specific genotypes are available in genetic surveys. Then, individuals can be classified by their *full categorical identities*, the combination of their identity covariate values for all identity covariates. For example, in a camera trap study, a full categorical identity might be “male, subadult” or “female, adult” or in a genetic study using 2 microsatellite loci, a full categorical identity would be the genotypes consisting of the number of DNA sequence repeats for each allele, where one individual’s full categorical identity might be “(144, 153), (134, 134)”.

A full categorical identity does not provide enough information to produce a unique identity, but it reduces the uncertainty in individual identity in two ways. First, it introduces deterministic constraints, preventing samples with conflicting covariate values from being allocated to the same individuals (e.g., a male and a female sample cannot be combined). Second, the number of categorical identity covariates, the number of category levels for each covariate, and the population frequencies of each category level, estimated by the model, inform the probability that any two samples that match at all identity covariates belong to a single individual. For example, two “male, subadult” samples more likely belong to the same individual than two male samples of unknown age class because the age class category subdivides the male category. Further, in a population with an uneven sex ratio (e.g. more males than females), the probability two male samples came from the same individual is lower than the probability that two female samples came from the same individual and vice versa if there are more females than males. Similarly, Augustine *et al.* (2018) demonstrate that the categorical SPIM offers a continuous model for sample uniqueness in genetic capture recapture studies, where the addition of microsatellite loci decrease the uncertainty in individual identity until no uncertainty remains if enough loci are available. Thus, the categorical SPIM provides an alternative to using the binary probability of uniqueness criteria (P(ID) and P(sib) Waits *et al.*, 2001) that guarantee the near certain uniqueness of genotypes and relaxes the requirement that genotypes are unique in the population.

Combining the features of SMR with those of the categorical SPIM would improve inference in typical SMR studies where categorical identity covariates may be available, and in categorical SPIM studies where it may be possible to classify individuals as marked, whether by natural or researcher-applied marks. Combining these the features of these two models will have the greatest utility in scenarios where each of these models alone do not produce unbiased and precise density estimates. As we will demonstrate, SMR can produce biased and imprecise density estimates when there are few marked individuals and/or data are sparse, whether due to a low abundance of animals in the population or a low baseline detection rate or detection probability. Categorical SPIM also produces biased and imprecise density estimates when data are sparse and the available number of identity categories is small (Augustine *et al.*, 2018). Further, both SMR and categorical SPIM have difficulty accommodating individual heterogeneity in detection function parameters, especially *σ*, which can introduce positive bias in abundance and density estimates (Augustine *et al.*, 2018). Augustine *et al.* (2018) demonstrated that this positive bias can be removed with enough categorical identity covariates in the categorical SPIM, so adding these to SMR should allow for improved density estimation in the presence of individual heterogeneity in detection function parameters. Similarly, the extra information stemming from the marked individuals should improve density estimation over the categorical SPIM in such scenarios with individual heterogeneity in detection function parameters, especially if the marking process is explicitly modeled and/or if telemetry data are available.

Here, we develop a model that combines the marked individual features of SMR, the possibility of explicitly modeling the marking process as provided by generalized SMR, the use of individual-linked telemetry data, and the categorical identity covariates of categorical SPIM. We also introduce the possibility that detection function parameters vary by the values of the categorical identity covariates, an idea introduced by (Augustine *et al.*, 2018), but not yet implemented. Finally, we describe an efficient MCMC algorithm to fit conventional, generalized, and categorical SMR using the individual by trap capture histories rather than the individual by trap by occasion capture histories used to date (e.g., Sollmann *et al.*, 2013; Whittington *et al.*, 2016). We then investigate the performance of the categorical SMR model via simulation and apply the model to a fisher (*Pekania pennanti*) data set where categorical identity covariates are available in the form of microsatellite loci, where the marking process allows for perfect individual identification, and where individual-linked telemetry data are available.

## 2 Methods – Data and Model Description

### 2.1 Methods – Spatial Mark-Resight Foundation

In the spatial mark-resight study design, a subset of the population is marked either by researcher-applied or natural marks. Then, the population is subjected to a resighting process where the individual identities of marked individuals may or may not be observed and the individual identities of unmarked individuals are not observable. Sightings of marked individuals whose identities can be observed are deterministically associated into a capture history as they are in a typical SCR model, but there are three types of latent identity samples specific to mark-resight – marked but unknown identity, unmarked, and unknown marked status samples. If there are no marked but unknown identity or unknown marked status samples, the true capture history is partially latent, with the marked individual component being fully observed. However, if either or both of these two sample types that were or could have been produced by marked individuals are observed, the true capture history is fully latent. The inferential process of SMR is then twofold. First, the latent identity samples are probabilistically associated with the individuals that could have produced them in the fully or partially latent true capture history; however, unlike unmarked SCR, these prob-abilistic associations are subject to the deterministic constraints provided by the observed sample types. Second, the probabilistically-resolved true capture history is used to estimate abundance and density while accounting for the uncertainty in individual identification.

Spatial mark-resight is exactly a typical SCR model with an altered observation model. Therefore, it uses the same process model for the allocation of activity centers across the landscape. We assume *N* activity centers, ***s***_1_…, *s_N_*, are distributed uniformly across a 2-dimensional state-space, e.g., ***s***_*i*_ ∼Uniform(𝓢), where 𝓢 is the 2-dimensional state-space. These activity centers are organized in the *N* × 2 matrix ***S***. Note, this specification implies that the marked and unmarked individual activity centers have the same spatial distribution, which is reasonable if animals are marked randomly with respect to space as should be the case for natural marks or a completely random marking process. If the marking process is not random with respect to space, the capture-recapture data generated from the marking process can be included in the model leading to a generalized SMR model (Whittington *et al.*, 2016). We will consider both the case of conventional SMR where individuals are resighted at *J^S^* spatial locations, *X^S^*, over *K^S^* occasions and generalized SMR where they are first captured and marked at *J^M^* spatial locations, *X^M^*, over *K^M^* occasions followed by the same sighting process as conventional SMR. The dimension of the matrices of spatial locations are *J^M^* × 2 and *J^S^* × 2, respectively, for *X^M^* and *X^S^*. Below, we will describe generalized SMR, with the implication that our description of the sighting process fully describes the observation model of conventional SMR.

Both the marking and sighting processes of generalized SMR are typical SCR observation models, with the capture history being partially or fully latent for the sighting process. We will consider both a Poisson and Bernoulli observation process for both observation components. For the Poisson observation model, we assume the observed counts for individual *i* at trap *j* summed over the *K^m^* capture occasions for observation type *m* are distributed following 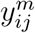 ∼ Poisson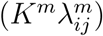 for *m* ∈ (M,S). We assume the expected counts for each individual at each trap on each occasion decline as an decreasing function of distance following 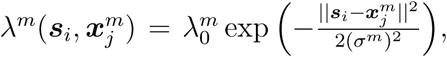 where 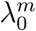 is the expected number of counts for a trap located at distance 0 from an activity center for observation type *m*, and *σ^m^* is the spatial scale parameter determining how quickly the expected counts decline with distance.

For the binomial observation model, we assume the detections for individual *i* at trap *j* summed over the *K^m^* capture occasions for observation type *m* are distributed following 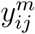∼ Binomial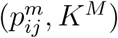 where 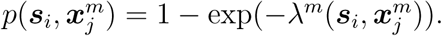 For convenience, we will refer to both the trap-level detections and trap-level counts as “counts” below.

The observed data consists of the fully observed marking process capture history for the *n^M^* marked individuals, the complete or partially-observed sighting process capture history for the marked individuals, and the sighting process capture histories for the latent identity samples. We denote the *n^M^* × *J^M^* capture history for the marking process as ***Y**^M^* and the *n^M^* × *J^S^* capture history for the sighting process of the marked individuals as ***Y***^*S.M*^. The ***Y***^*S.M*^ matrix is the marked individual submatrix of the true sighting history ***Y**^S^*. ***Y**^S^* is the *n^cap^* × *J* complete sighting history which we wish to probabilistically reconstruct, where *n^cap^* is the total number of individuals captured, both marked and unmarked. The marked individual sighting history, ***Y***^*S.M*^, is fully observed unless there are some samples where marked status can be determined but individual identity cannot, or if there are unknown marked status samples that belong to marked individuals. In fact, this matrix could possibly consist of all zero entries if the individual identity could not be determined for any of the marked status samples and thus, it is completely latent as we will consider in the fisher application.

The observed counts of each of the three latent identity sighting sample types can be aggregated to trap by occasion counts and stored in separate *J^S^* × *K^S^* dimension matrices for the marked but no identity, unmarked, and unknown marked status samples (Whittington *et al.*, 2016), or if counts are summed over occasions as we assume, they can be aggregated to trap level counts and stored in *J^S^* length vectors. However, because we will be associating categorical identities with each sample, the counts cannot be aggregated. The counts could be summarized by their traps of capture; however, to ease updating the latent data, we chose to store each count in a matrix row indicating the trap of capture. More specifically, let *n^M.unk^*, *n^um^*, and *n^unk^* be the number of marked but no identity, unmarked, and unknown marked status samples, respectively. Then ***Y**^S.M.unk^* is the *n^M.unk^* × *J^S^* matrix of marked but no identity samples, ***Y**^S.um^* is the *n^um^* × *J^S^* matrix of unmarked samples, and ***Y**^S.unk^* is the *n^unk^* × *J^S^* matrix of unknown marked status samples. These latent identity capture histories consist of all zero entries except for a single “1” entry indicating the trap of capture.

To visualize the relationship between the true and observed sighting histories, we provide an example of hypothetical true and disaggregated sighting histories for two unmarked individuals captured at *J* =3 traps. The true but unobserved unmarked individual capture histories are shown in *Y^S^*, where the first individual was captured once in trap 1 and once in trap 3, and the second individual was captured twice in trap 1, once in trap 2, and once in trap 3. In this example, the marked status (unmarked) was observed for all but the 2^nd^ individual’s captures in traps 2 and 3. Thus, the first individual’s captures are disaggregated into the first two rows of ***Y**^S.um^* the second individual’s captures in trap 1 are disaggregated into the second two rows of ***Y**^S.um^* and the second individual’s captures in traps 2 and 3 are disaggregated into ***Y**^S.unk^*.

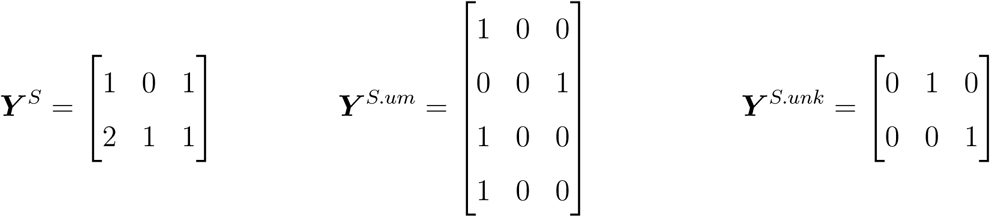

The latent identity marked individual samples are similarly disaggregated into ***Y**^S.M.unk^* if their marked status (marked) is determined and into ***Y**^S.unk^* if their marked status cannot be determined, along with the unknown marked status samples stemming from the unmarked individuals. The inferential process is then to probabilistically reassemble these latent identity samples, given the deterministic constraints introduced by the observed sample types, into the latent components of ***Y**^S.true^*, the augmented (described below), partially latent true sighting history, and then estimate abundance and density, incorporating this uncertainty in individual identity.

### 2.2 Methods – Categorical Spatial Mark-Resight

Here, we extend SMR to incorporate the class structured individual identity model of Augustine *et al.* (2018) where class membership is determined by an individual’s set of values for *n_cat_* categorical covariates, which we will call an individual’s *full categorical identity*, and where multiple individuals in the population may share the same full categorical identity. The joint density and categorical identity process model is exactly the same as the unmarked SCR-based model described in Augustine *et al.* (2018), while the observation model introduces the possibility of dividing the population into marked and unmarked individuals, with complete identities possibly available for the marked individuals.

Associated with the individual activity centers is an *N* × *n^cat^* categorical identity matrix ***G***, where ***g**_i_* is the full categorical identity of the individual having activity center ***s**_i_.* Each of the *l* categorical covariates have 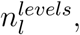 which we assume are all known and indexed sequentially by 1,…, 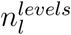(Wright *et al.*, 2009). The values of ***G*** are then determined by the population category level probabilities for each covariate. More specifically, the category level probabilities for covariate *l* are *γ_l_*, each of length 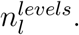 We then assume the categorical identity of each individual for each covariate are distributed following the covariate-specific category level probabilities following *g_il_* ∼ Categorical(*γ_l_*) and that category levels are distributed independently across identity covariates and individuals.

The partially latent observation process is the same as typical SMR; however, we consider that detection function parameters may depend on the levels of categorical identity covariates. Theoretically, any categorical feature observable or partially observable in an SMR survey that may determine the value of detection function parameters, such as individual sex or age class, can also be used as a categorical identity covariate. We consider the simplest case where the detection function parameters vary by only one categorical identity covariate; however, detection function parameters may vary by multiple categorical identity covariates using additive effects on the logit scale. We define the first categorical identity covariate, ***g*.**_1_, to be the identity covariate that determines the 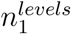 detection function 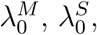 and *σ* parameters, now redefined to be vectors of length 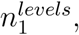 e.g., 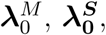 and ***σ***. Further, the population-level frequencies of these category levels are already stored in ***γ***_1_ described above, which can be used to update missing covariate values. For example, ***g*.**_1_ may be the sexes of the *N* individuals in the population in which case, 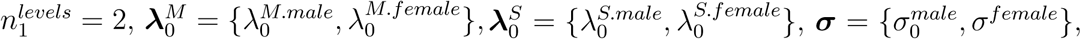 and *γ*_1_ is a vector of length 2 with elements P[sex_*i*_]=male and P[sex_*i*_]=female. These categorical covariate-specific detection functions allow for individual heterogeneity in detection function parameters to be reduced, which has been shown to introduce positive bias in density estimates from latent identity SCR models when not modeled (Augustine *et al.*, 2018). Further, the category level probabilities will not be estimated correctly if detection probability differs across category levels, a problem resolved by the use of covariate-specific detection functions.

The observed data in the unmarked SCR-based categorical SPIM (Augustine *et al.*, 2018) consists of two linked data structures. First, ***Y**^obs^* is a capture history matrix with one row per sample, indicating the trap at which each latent identity sample was recorded with a “1” with all other entries being “0”. Second, ***G**^obs^* is the matrix of observed categorical covariate(s), where each row stores the observed category levels of the latent identity samples stored in the same row of ***Y**^obs^*. These category levels are indexed sequentially as described above or recorded as a 0 if not observed. Above, we split the observed latent identity counts by sample type, so we will need to split ***G**^obs^* by sample type: ***G**^M.unk^* is the *n^M.unk^* × *n^cat^* categorical identity matrix for the marked but unidentified samples, ***G**^um^* is the *n^um^* × *n^cat^* categorical identity matrix for the unmarked samples, and ***G**^unk^* is the *n^unk^* × *n^cat^* categorical identity matrix for the unknown marked status samples. Unlike the categorical SPIM, the full categorical identities for a subset of the individuals, specifically, those that are marked, are partially (if missing covariate values) or fully known and stored in ***G**^M^* of dimension *n^M^* × *n^cat^* and linked to ***Y**^M^* and ***Y**^S.M^* by the *i* dimension.

Using the hypothetical true and observed sighting histories for two unmarked individuals above, we suppose individual 1 has full categorical identity “3, 6, 2” and individual 2 has full categorical identity “1, 2, 2”. In the example above, the marked status for the first individual’s 2 samples was determined, as were the second individual’s captures in trap 1, so their observed categorical identities are stored in ***G**^S.um^*. Then, the marked status of the second individual’s captures in traps 2 and 3 was not determined, so the observed categorical identities are stored in ***G**^S.unk^*. In this example, the value of each categorical identity covariate was always observed; however, we allow missing covariate values coded with a “0”.

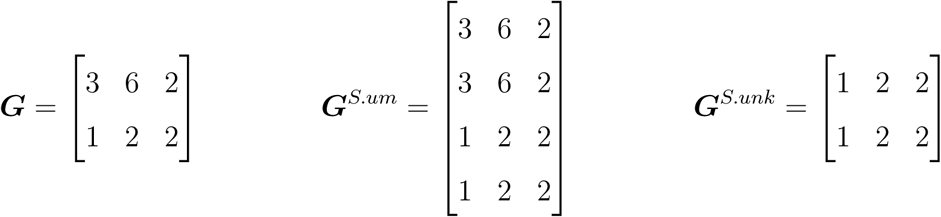

The inferential challenge is then to probabilistically resolve the individual identities of the latent identity samples, subject to the deterministic sample type and categorical identity constraints, and estimate abundance and density incorporating this uncertainty in individual identity. This is accomplished using data augmentation (Royle *et al.*, 2007). Both ***Y**^M^* and ***Y**^S.M^* are augmented up to dimension *M* × *J^M^* and *M* × *J^S^*, respectively, with *M* chosen to be much larger than the expected abundance. We will rename the augmented sighting history ***Y**^S.true^* because it will be allocated latent identity samples from both the marked and unmarked components of the population in order to probabilistically reconstruct the complete, true sighting history. The augmented individuals have fully unobserved full categorical identities which requires ***G**^M^* to be augmented up to dimension *M* × *n^cat^.* We will rename the augmented categorical identity matrix ***G**^true^* to match ***Y**^S.true^*.

Following Royle *et al.* (2007), the vector ***z*** is introduced to indicate that individuals are in the population with value “1” and “0” otherwise, with the first *n^M^* individuals’ values fixed to “1” because they are known to be in the population. Then, it is assumed that *z_i_* ∼ Bernoulli(*ϕ*) inducing the relationship *N* ∼ Binomial(*M*, *ϕ*). Population abundance, 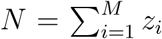 is a derived parameter and population density, *D*, is 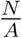 where *A* is the state-space area. Finally, the identity to which each latent identity sample is assigned to on each MCMC iteration is a derived parameter, all of which are stored in vector, ***ID***, of size *n^latent^*, the total number of latent identity samples of all types. The posteriors for *ID* can be post-processed to calculate the pairwise posterior probabilities that any two samples were produced by the same individual. The algorithms for updating all parameters, including those for updating ***Y**^S.true^* subject to the deterministic constraints stemming from the sample types and the observed categorical covariates can be found in Appendix A.

## 3 Simulations

### 3.1 Simulation Specifications

We conducted simulations across eight scenarios to illustrate two main features of SMR and categorical SMR – 1) that conventional SMR estimates with sparse data and/or with few marked individuals produce biased and imprecise estimates, similar to Augustine *et al.* (2018) that demonstrated unmarked SCR produces unbiased and imprecise estimates when data are sparse, and 2) that bias can be removed and precision increased by introducing categorical identity covariates. We first replicated scenarios A1-A4 from Augustine *et al.* (2018) with the caveat that we considered that 10% of the population was marked. These four scenarios constituted a factorial design with *D* ∈ (0.17,0.35), corresponding to *N* ∈ (39, 78), and *σ* ∈ (0.5,1). For scenarios with *σ* = 0.5, we set 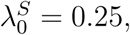 which led to an expected number of captures per individual of 1.65. To maintain a similar level of data sparsity in scenarios with *σ* = 1, we set 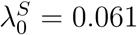 to maintain the same 1.65 expected captures per individual. To specify approximately 10% of the population to be marked, we used 4 marked individuals when *N* = 39 and 8 marked individuals when *N* = 78.

We then considered four additional scenarios (A1b - A4b) where data were more sparse, but more marked individuals were available to demonstrate that the increased precision stemming from additional marked individuals can be offset by increasing data sparsity. For scenarios with *σ* = 0.5, we lowered 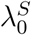 to 0.125, leading to an expected number of captures per individual of 0.8 (an average including the 0 values of uncaptured individuals). These scenarios are closer to the estimated mean number of captures per individual (samples/*N̂*) observed in Sollmann *et al.* (2013, 1.08), Kane *et al.* (2015, 0.86), and the most data sparse application of Rich *et al.* (2014, 0.94). Then for scenarios with *σ* = 1, we lowered 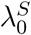 to 0.03, matching the 0.8 expected number of captures per individual. To specify approximately 20% of the population to be marked, we used 8 marked individuals when *N* = 39 and 16 marked individuals when *N* = 78. For all eight of these scenarios, we fit a regular conventional SMR model, followed by the conventional version of the categorical SMR model with 1-8, 2-level categorical identity covariates added sequentially, leading to the unique number of identity categories increasing with base 2 (2, 4, 8, 16, 32, 64, 128, 256). We assumed there were no marked but unidentifiable or unknown marked status samples, leaving unmarked samples as the only latent identity sample type.

All simulations were conducted on a 9 × 9 trapping array with unit spacing. The state-space was defined by buffering the trapping array by 3 units. Individuals were subjected to captured over *K* =10 occasions. For all 8 scenarios, we simulated 125 data sets and fit the conventional SMR or categorical SMR model, using the posterior modes for point estimates and the highest posterior density intervals (HPD) for interval estimators. We computed the bias and mean squared error (MSE) of the point estimates and the 95% coverage and 95% HPD interval widths of the interval estimators to quantify frequentist bias and coverage, and estimator accuracy and precision. Finally, we estimated *n_um_*, the number of unmarked individuals that produced the unmarked samples to compare to the simulated values and to explore the influence of the uncertainty in *n_um_* on the precision of the density estimates.

### 3.2 Simulation Results

The conventional SMR estimates were approximately unbiased for the small *σ* scenarios A1 and A2 (Table B1) and positively biased for the larger *σ* scenarios, by 7% for the high density scenario A3 and by 22% for the low density scenario A4. In both of these large *σ* scenarios, the bias was largely removed once 4 identity categories were available. For the scenarios with more sparse data and more marked individuals (A1b-A4b), there was about 5-7% positive bias in the higher density scenarios A1c and A3c with about 20% positive bias in the lower density scenarios A2c and A4c. This bias was effectively removed by adding identity categories (Figures 1 and 2, Table B1), but the number required for each scenario varied, with more being required for the larger *σ* scenarios A3c and A4c. The frequentist coverage for the SMR and categorical SMR estimators was close to nominal, on average, for all scenarios (Table B1). Estimator accuracy and precision were improved by adding identity categories in all scenarios, but generally with diminishing returns (Figures 1 and 2, Table B1). Two exceptions to this trend are found in scenarios A3b and A4b (Figure 2), with more sigmoidal relationships between the number of identity categories and precision where adding 2 identity categories did not offer much improvement, but using 4-16 offered greater improvements before precision and accuracy gains tapered off again. The density estimates in scenarios A1c-A4c with more marked individuals, but fewer captures per individual were less precise and accurate than scenarios A1-A4 with fewer marked individuals and more captures per individual, requiring substantially more identity categories to achieve the same precision and accuracy (Figures 1 and 2, Table B1).

**Figure 1:**
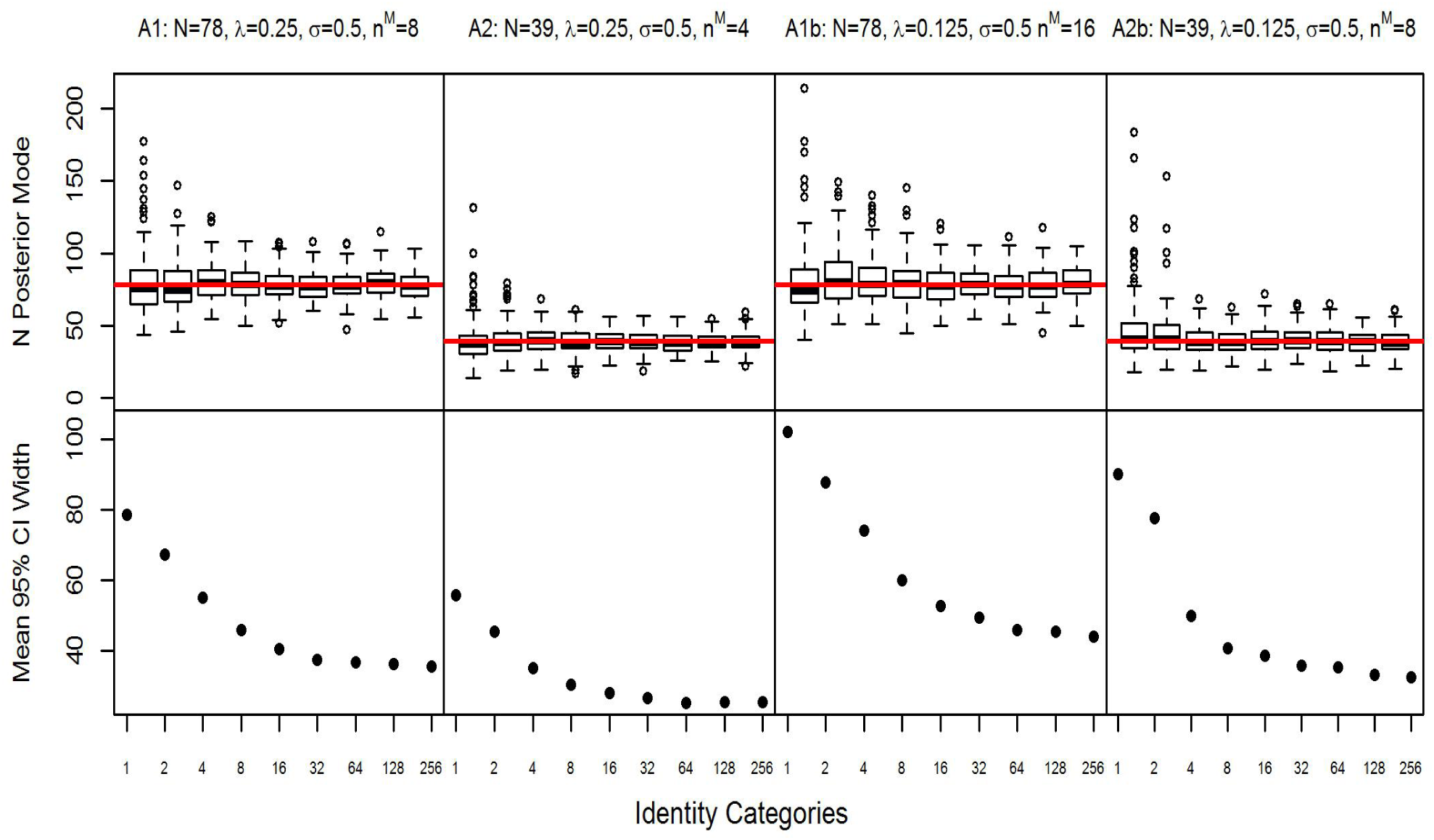
Posterior modes and mean 95% credible interval width of abundance (N) estimates plotted against the number of identity categories in Scenarios A1, A2, A1b, and A2b with detection function spatial scale parameter *σ* = 0.5 and baseline detection rate λ_0_ = 0.25 with 10% of the population marked and λ_0_ = 0.125 with 20% of the population marked. Note the number of identity categories increase exponentially.

**Figure 2:**
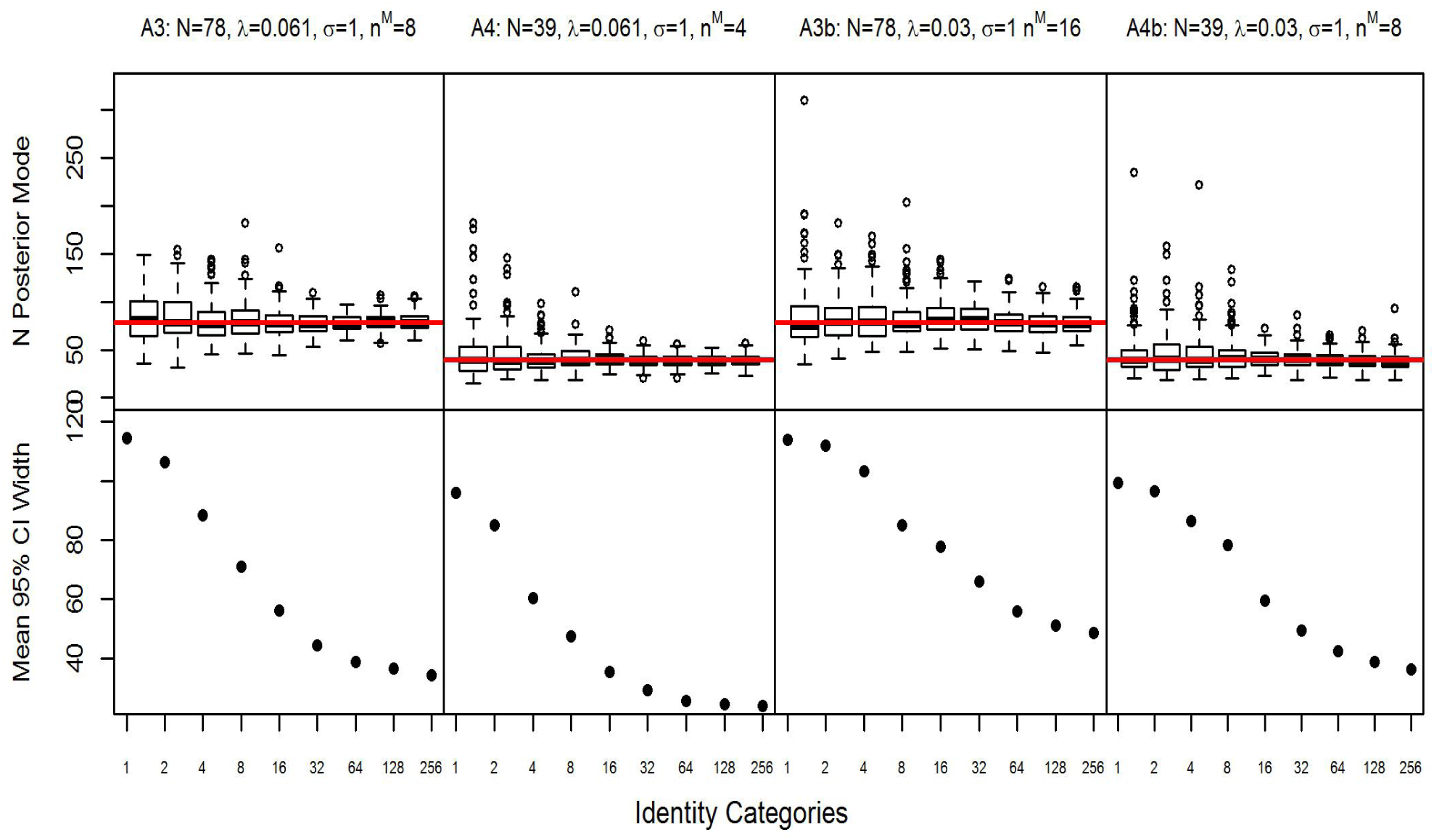
Posterior modes and mean 95% credible interval width of abundance (N) estimates plotted against the number of identity categories in Scenarios A3, A4, A3b, and A4b with detection function spatial scale parameter *σ* = 1 and baseline detection rate λ_0_ = 0.061 with 10% of the population marked and λ_0_ = 0.03 with 20% of the population marked. Note the number of identity categories increase exponentially.

Unlike the unmarked SCR and categorical SPIM simulations in Augustine *et al.* (2018), the bias in the sparse data SMR and categorical SMR estimates was always positive. Generally, the bias of the SMR estimates was of a similar magnitude as the unmarked SCR estimates; however the SMR estimates were more accurate and precise, with the SMR advantage over the unmarked estimator decreasing as the number of identity categories increased. Once 256 identity categories were available, the improvement in accuracy and precision of the SMR estimates over the unmarked SCR estimates was negligible. Similar to the unmarked SCR and categorical SPIM estimates, the SMR and categorical SMR models overestimated the number of latent identity individuals captured, *n^cap^* in Augustine *et al.* (2018) and *n^um^* in the present simulations, when no or few identity categories were available. Also similar is the fact that adding more identity categories reduced the uncertainty in *n^um^*, which is the mechanism reducing uncertainty in abundance and density.

## 4 Application – Conventional and Generalized SMR Analysis of Fisher Genetic Capture-Recapture

Similar to Augustine *et al.* (2018), we applied the proposed categorical SMR models to a genetic capture-recapture data set, which provided abundant categorical identity covariates in the form of microsatellite loci. A desirable feature of applying these models to microsatellite loci is that they provide enough categorical identity covariates to guarantee near certainty in individual identities when all loci are used (as determined by P(ID) and P(sib) criteria), which provides metrics to which scenarios using fewer loci can be compared. For example, when all observed genotypes in a sample can be considered unique with high probability, the resulting density estimate should be better than estimates using fewer loci that permit uncertainty in individual identity. The number of individuals captured, typically not known in SMR, is also known with near certainty when unique genotypes can be considered unique identities with near certainty. In contrast to Augustine *et al.* (2018), which used an unmarked model extended to allow categorical identity covariates, this data set provided several additional sources of information to improve the probabilistic association of latent identity samples, and thus density estimates. First, several previously captured individuals, marked with GPS collars or conceivably marked with other unique identifiers observable upon live capture (see below), were known to be in the population during the partial identity capture process. Second, telemetry data were available for a subset of these marked individuals. Third, capture-recapture data from the marking process was available so that the marking process, where individuals can be identified with certainty, could be explicitly modeled.

This data set is from a joint live-capture and hair snare survey of fishers in the province of Alberta, Canada (Stewart *et al.*, 2017, 2018), where individual sex and up to 15 microsatellite loci were available to use as categorical identity covariates. We used the 8 microsatellite loci that were amplified for all individuals, which included markers Ggu101, Lut604, Ma-1, Ma-19, MP144, MP182, and Mvis72 (previously identified by Jordan *et al.*, 2007; Davis & Strobeck, 1998; Dallas & Piertney, 1998; Fleming *et al.*, 1999; Duffy *et al.*, 1998) which produced 10, 10, 10, 5, 13, 16, 11, and 14 unique loci-level genotypes, respectively. Live capture occurred at 37 locations over 129 days with individual trap sites being used for a mean of 46 days (range from 2-115). During this time, 24 fishers were captured (8 male, 16 female) from 1 to 4 times each (mean of 1.5). GPS collars were placed on 10 individuals (5 male, 5 female), which allowed for individuals to be identified upon recapture. In the original study, the individual identities of the 14 individuals that were not fitted with a GPS collar were determined by their 8 loci genotypes. For the purposes of this application, we assume that all 24 live-captured individuals were marked in a manner that allowed them to be identified with certainty upon recapture during the marking process (e.g., with collars, passive integrated transponder tags, or ear tags) because we do not assume that unique genotypes correspond to unique individuals with certainty due to the possibility of the shadow effect (where more than one captured individual share the same genotype; Mills *et al.*, 2000).

Hair snares were deployed in a stratified random design across 64 4 × 4 km grid cells. Each hair snare was baited with a 5 kg piece of beaver carcass and lure (O’Goreman’s Long Distance Call). These baits were applied to the sample tree, which was wrapped in barbed wire to obtain a hair sample as a fisher climbed the tree to access the bait (sensu Stewart *et al.*, 2018). Sites were checked monthly, when bait was replenished, hair samples were collected, and barbed wire was torched to eliminate the possibility of DNA contamination between capture occasions. This led to four monthly capture occasions. Assuming a unique 8 loci genotype is unique in the population with high probability (justified by the probability of identity criteria), twenty-four individuals (10 male, 14 female) were detected via hair snares from 1 to 8 times (mean of 2.4); however, some of these individuals were not live captured and vice versa. A total of 57 hair snare samples providing 8 loci genotypes and 25 hair snare samples providing fewer than 8 loci genotypes were collected. The partial genotype samples did not provide the sex marker, but provided from 1-6 microsatellite loci each (mean 1.6). The spatial distribution of captures for both capture processes are illustrated in Figure 3.

**Figure 3:**
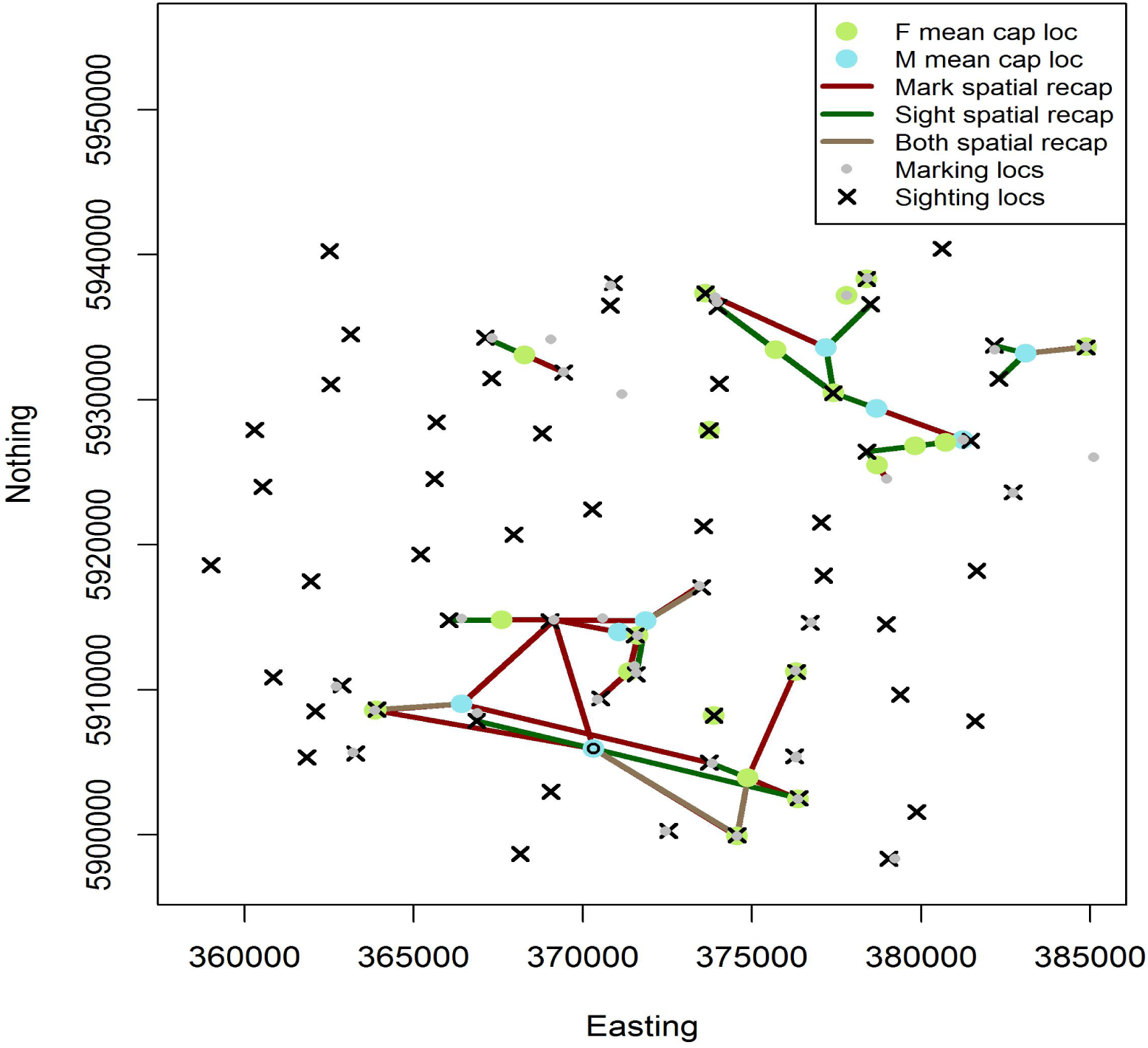
The spatial configuration of marking (live capture) and sighting (hair snare) locations, mean marking/sighting locations and spatial recaptures for the fisher application. The male marked with a circle has an especially wide distribution of spatial recaptures, especially during the sighting (hair snare) survey.

This joint live-capture and hair snare data set fits within a generalized SMR framework in the following way. First, we will ignore the genotypes. The live-capture survey constitutes a *marking process* where all individual identities can be determined. Then, the hair snare survey constitutes a *resighting process* with the caveat that without the certain identities provided by genotypes, the sightings for the marked individuals cannot be deterministically linked. Therefore, the true sighting histories for the marked individuals are fully latent and must be treated as *unknown marked status* samples. Telemetry data are also available during the hair snare survey (sighting process) to improve the estimates of *σ* and the activity centers of the telemetered individuals. If the live-capture data are discarded, this data set fits within the framework of conventional SMR, where it is known that a certain number of marked individuals exist in the population during resighting, and the telemetry data provides information about where their activity centers are located. However, in this application, live traps were targeted towards areas with higher levels of fisher sign (hair samples or photo events from remote cameras), while hair snares were placed more widely, suggesting that conventional SMR will underestimate density for this data set and that generalized SMR will be necessary for proper inference once complete individual identities are discarded, precluding a typical SCR analysis.

From either the conventional or generalized SMR foundation, we can introduce the microsatellite loci as categorical identity covariates, transitioning to a categorical SMR framework. In this example, there were 57 sighting process (hair snare) samples that provided 9 categorical identity covariates (sex plus 8 microsatellite loci, partial genotype samples excluded). We analyzed this data set in three ways, sequentially adding microsatellite loci until there was no remaining uncertainty in individual identity in each analysis. The three analyses were 1) only using the sighting (hair snare) observations in a conventional SMR model, 2) using the sighting (hair snare) observations and telemetry data in a conventional SMR model, and 3) using both the marking (live-capture) and sighting data (hair snare) in a generalized SMR model without the telemetry data. Microsatellite loci were added in the order of least to most diverse so that the lowest level of categorical identity information was available for each *X*-loci estimate. Due to expected sex differences is fisher space-use, we used identity covariate-specific detection functions for the sex covariate to reduce individual heterogeneity in detection function parameters. In all three analysis types, we included a scenario where the 25 partial genotype samples were added to the full 8 loci data set.

For all models, MCMC chains were run for 100,000 iterations, with the first 5,000 discarded, and thinned by 20. Posterior modes were used for point estimates and the highest posterior density (HPD) intervals were used for interval estimates. To provide benchmarks to compare to the conventional and generalized categorical SMR estimates, we fit regular SCR models to both the marking (live capture) and sighting (hair snare) process data assuming unique 8 loci genotypes were unique individuals (justified by the P(sib) criterion). We refer to these estimates as the “marking SCR estimate” and “sighting SCR estimate” below. The 8 loci genotypes also allowed us to know the number of unique individuals seen during the sighting process, *n^unk^*, with near certainty (24), a quantity that is generally not known in SMR.

### 4.1 Application – Results

The conventional SMR density estimates with and without telemetry (Table 1) were generally lower than the reference sighting SCR estimates. This pattern is consistent with the expected negative bias when applying conventional SMR to a population where the marked individuals are not a random sample of individuals from the state-space. The density estimates with and without telemetry stabilized after at least 3 loci were used, with the 1-3 loci estimates exhibiting less bias (relative the reference SCR estimates) when telemetry was used. The precision of the density estimates was similar for the 4-8 loci estimates with and without telemetry, which showed a slight improvement in precision over the 3 loci estimates. The 3-8 loci conventional SMR models correctly estimated the number of individuals producing the latent identity samples, *n^unk^*, except for the 6 loci estimate without telemetry. The 6 loci model estimated *n^unk^* to be 25, erroneously splitting the samples for one male with spatial recaptures distributed over a particularly large area (see Figure 3) across two individuals. When this individual’s samples were erroneously split, the male *σ* estimate was substantially lower than the models correctly estimating *n^unk^* to be 24, indicating strong individual heterogeneity in *σ*.

**Table 1:**
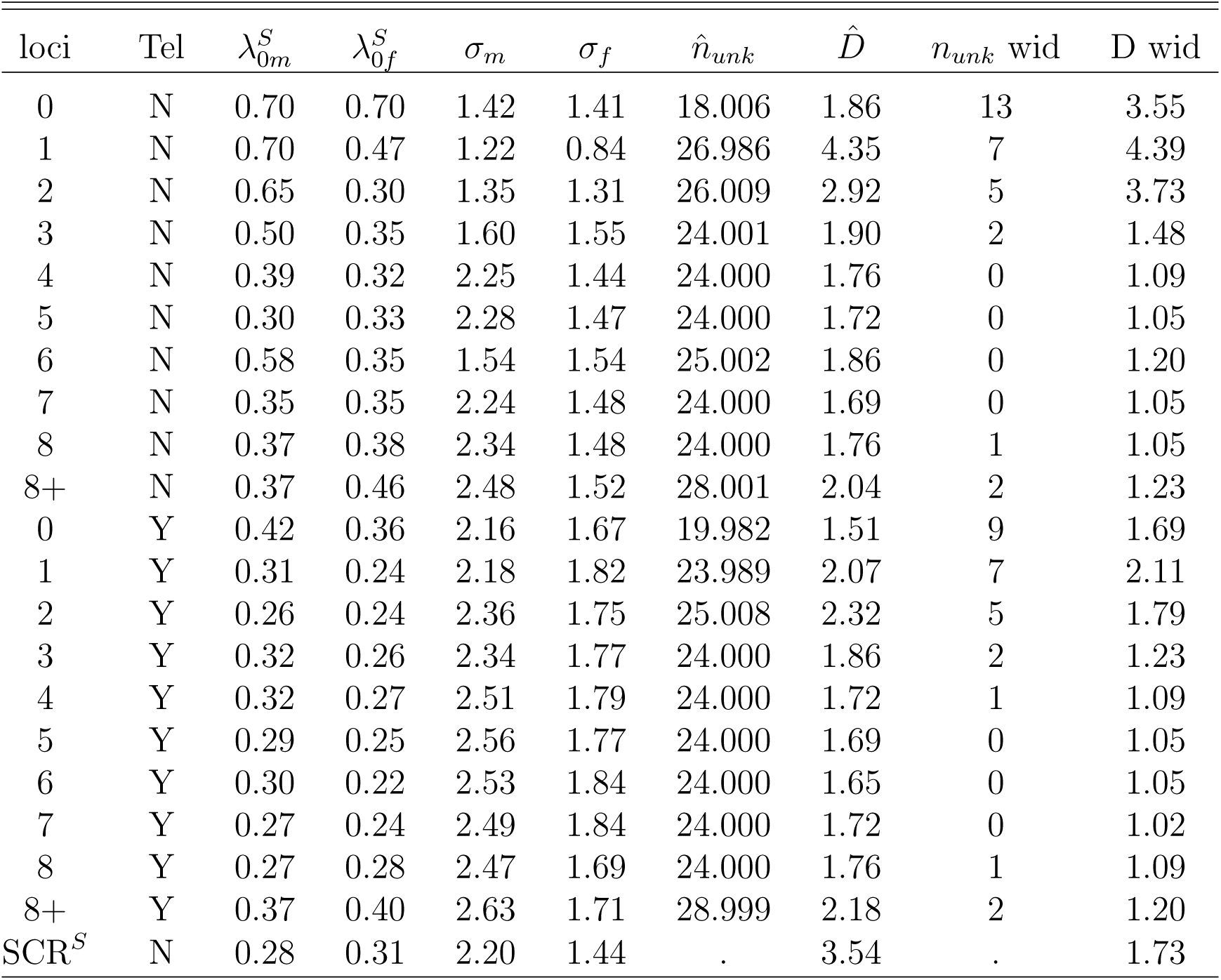
Conventional spatial mark-resight estimates with and without the use of telemetry data (Tel) for the fisher data set using 0-8 loci and including partial genotypes into the 8 loci analysis. SCR^*S*^ is the complete data SCR estimate for the sighting data set. Columns include the sex-specific λ and *σ* parameter estimates, followed by the estimated number of individuals that produced the partial identity samples, the estimated density, and the mean 95% credible interval widths for the latter two quantities.

**Table 2:**
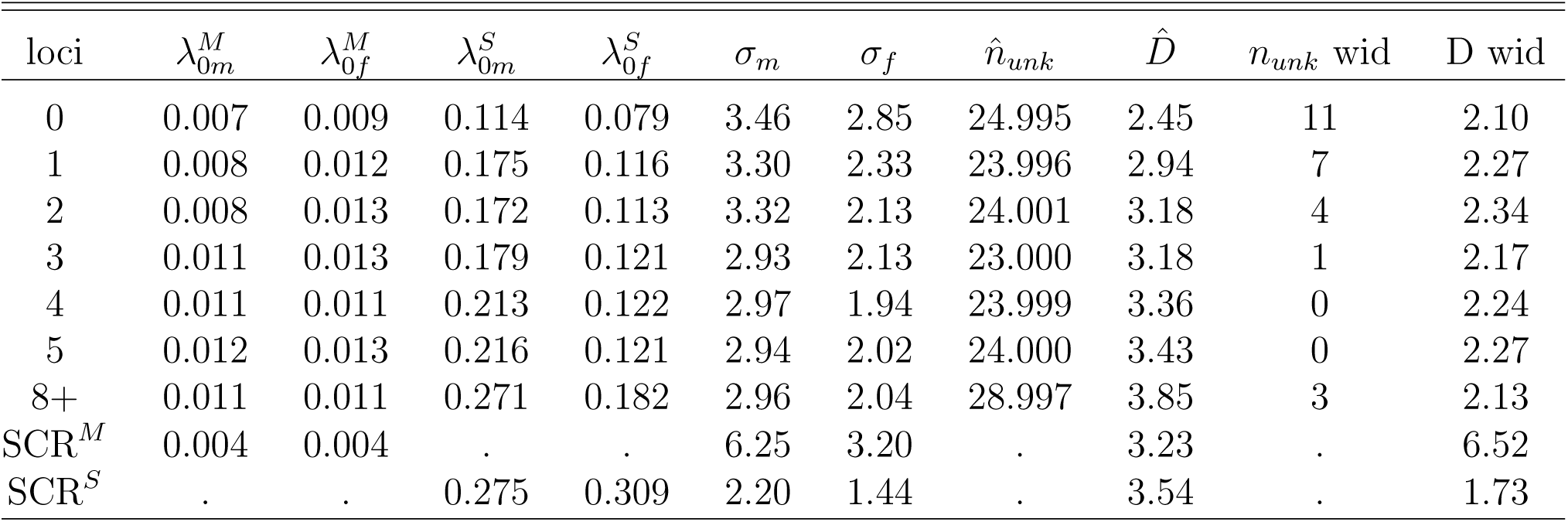
Generalized spatial mark-resight estimates for the fisher data set using 0-5 loci and including partial genotypes into the 8 loci analysis. SCR^*M*^ and SCR^*S*^ are the complete data SCR estimates for the marking and sighting data sets, respectively. Columns include the sex-specific marking and sighting λ parameter estimates and the sex specific *σ* estimates, followed by the estimated number of individuals that produced the partial identity samples, the estimated density, and the mean 95% credible interval widths for the latter two quantities.

While the conventional SMR density estimates appeared to be negatively biased, the *σ* estimates for the 3, 4, 5, 7, and 8 loci conventional SMR estimates without telemetry were very similar to the sighting SCR estimates. The *σ* estimates for the 3-8 loci estimates using telemetry were slightly higher than the sighting SCR estimates; however, these conventional SMR *σ* estimates should not be expected to match the sighting SCR analysis exactly because the latter did not use the telemetry data. Adding the 25 partial genotype samples slightly increased the density point estimates and slightly decreased the precision of the density estimates in the conventional SMR estimates with and without telemetry. The model without telemetry estimated that these 25 samples belonged to 4 individuals, while the model using telemetry estimated that they belonged to 5 individuals, likely due to the larger *σ* estimate of this model allowing the partial identity samples to be allocated to individuals with activity centers further from the traps where samples were recorded. Of these two models, the one with telemetry, which estimated 1 additional captured individual than the model without telemetry, saw a larger increase in the density estimate over the 8 loci estimate without partial genotypes.

The generalized SMR density estimates were higher than the conventional SMR estimates, and similar to both the marking and sighting SCR density estimates. This result is consistent with the removal of negative bias by modeling the marking process explicitly Whittington *et al.* (2016). The generalized SMR density estimates were less precise than the conventional SMR density estimates, which can be partially explained by the fact that if conventional SMR estimates are negatively biased due to a non-random spatial distribution of the marked individuals, the variance will be underestimated. The generalized SMR density estimates were more precise than the marking SCR density estimate, but less precise than the sighting SCR density estimate, which reflects the fact that some of the uncertainty in the marking process density estimate propagated into the uncertainty in the generalized SMR density estimate. The generalized SMR *σ* estimates were a weighted average of the marking and sighting SCR *σ* estimates, with the estimates being closer to the sighting SCR estimates because the average number of captures per individual was higher during the sighting process. The marking process λ_0_ estimates were higher in the generalized SMR model than the marking process SCR model due to the lower *σ* estimates in the generalized SMR model. Similarly, the sighting process λ_0_ estimates were lower in the generalized SMR model than the sighting process SCR model due to the higher *σ* estimates in the generalized SMR model.

The generalized SMR density estimates stabilized after 2 loci were used, with no uncertainty left in individual identity once 5 loci were used and the precision of the density estimates was similar for the 0-5 loci estimates. The 0 and 1 loci estimates were 28% and 14% lower than the 5 loci estimate, but were similarly precise. The 3 loci estimate incorrectly estimated that the latent identity samples belonged to *n^unk^* = 23 individuals, one too few, but the density estimate was not noticeably affected by this slight underestimation. The density estimate using the 25 partial genotypes was similar to the 2-5 loci estimates, with similar precision. Like the conventional SMR estimate using telemetry, the generalized SMR model estimated that the 25 partial genotype samples belonged to 5 individuals.

### 4.2 Application – Discussion

The conventional SMR fisher estimates generally behaved similarly to those seen in the conventional SMR simulations. Density estimates were biased when using few identity categories, and this bias was removed with the addition of more identity categories. Unlike the categorical SMR simulations; however, the fisher categorical SMR density estimates using the fewest identity categories (2 sex categories, 0 loci) were negatively biased, switching to positive bias with the addition of 1 and 2 loci. The bias seen in the categorical SMR simulations with fewer identity categories was always positive. Both negative and positive bias with fewer identity categories was observed in the categorical SPIM estimator (Augustine *et al*., 2018); therefore, we suspect either positive or negative bias is possible when applying SMR and categorical SMR to sparse data sets with few marked individuals, depending on the exact simulation specifications.

As expected, the addition of telemetry data and an explicit model for the marking process improved SMR density estimation, especially when there were fewer identity categories. The conventional SMR density estimates using 0-3 loci were less biased (relative to the 4-8 loci estimates) when using telemetry data and this lower bias was associated with better estimates of *n^unk^* and *σ_m_.* The 0-3 loci generalized SMR density estimates were less biased than the conventional SMR estimates without telemetry, each relative to their 8 loci estimates, which was also associated with better estimates of *n^unk^* and *σ_m_.* Using the partial 25 genotype samples had a negligible effect on the categorical SMR density estimates; however, the majority of the partial genotypes were only observed at 1 locus, with the mean multilocus genotype being observed at 1.6 locus, so very little identity information was available. Further, it is possible the partial genotype samples contained genotyping errors that reduced their usefulness (see Augustine *et al.*, 2018, for more discussion of the application of categorical SPIM models to microsatellite genotypes).

Similar to the black bear application of Augustine *et al.* (2018), it appears that one particular male fisher (Figure 3) had an especially wide distribution of spatial recaptures during the sighting process, perhaps introducing individual heterogeneity in *σ_m_.* When combining the marking and sighting processes, the spatial extent of this male’s spatial recaptures became less extreme relative to other males; however, individual heterogeneity likely still remained. Correctly connecting this male’s samples in the conventional SMR models, which required at least 3 loci, was influential to the magnitude of bias in the density estimate. In the conventional SMR model with fewer than 3 loci, this male’s samples were split between two individuals, leading to a large reduction in the *σ_m_* estimate. The larger *σ_m_* estimate observed when using either the telemetry data in the conventional SMR models or a model for the marking process in the generalized SMR models appeared to counteract the problems introduced by individual heterogeneity in *σ.*

It appears that explicitly modeling the marking process was necessary for unbiased density estimates in this fisher data set where animals were not marked randomly with respect to space (Whittington *et al*., 2016). One potential downside of modeling the marking process not previously discussed is that if the data from the marking process are sparse, as it was in the fisher data set, the precision of the density estimates will be eroded. Further, it is not clear that the *σ* parameters between the marking and sighting process were the same. The marking process SCR *σ* point estimates for males and females were larger than the sighting process SCR *σ* estimates, with little posterior overlap (data not shown). It is possible that fisher space use changes across the sampling season, as individuals may have been preparing for breeding in the final months of data collection. Alternatively, this difference could be a consequence of individual heterogeneity in *σ* for both sexes, with the larger *σ* individuals being more disproportionately observed in the marking process which had lower baseline detection rates than the sighting process. A third possibility is that activity centers are transient through time (Royle *et al.*, 2016) and the average duration a live capture (marking process) site was open was 2 months, while all hair snares (sighting process) were operational over 4 months, allowing individuals to move further distances. In general, the consistency of *σ* through space, time, and between observation methods should be treated with more scrutiny, and the repercussions of *σ* inconsistency should be investigated further (Popescu *et al.*, 2014; Tenan *et al.*, 2017).

## 5 Discussion

We combined the features of SMR and the categorical SPIM that improve the probabilistic association of latent identity samples and thus density estimation into the categorical SMR model. Specifically, the categorical SMR model allows for a subset of the individuals to be marked, individual-linked telemetry data for marked individuals, a capture history generated by the marking process, and any number of categorical identity covariates. Further, we developed one version of the categorical SMR model that allows for the detection function parameters to vary as a function of the category levels for one of the identity covariates, which will reduce the magnitude of unexplained individual heterogeneity in detection function parameters if this individual heterogeneity can be explained by one of the identity covariates, such as individual sex. This feature can be added to the categorical SPIM to improve inference when no marked individuals are available.

Our simulation study showed that similar to unmarked SCR (Augustine *et al.*, 2018), SMR density estimates can be biased and imprecise when data are sparse and/or there are few marked individuals. The simulation scenarios that matched the unmarked SCR scenarios of Augustine *et al.* (2018) showed a similar level of bias, but the marking of just 10% of the population greatly improved the precision of density estimates. The second set of scenarios we considered with 20% of the population marked and λ_0_ reduced by 50% demonstrated that gains in accuracy and precision from adding more marked individuals can be more than offset by increasing data sparsity. We demonstrated that the addition of categorical identity covariates can remove bias and increase precision in all of these scenarios, introducing another method to improve accuracy and precision in SMR models if categorical identity covariates are available.

The fisher application demonstrated that the addition of telemetry data improved the probabilistic identity associations in conventional SMR as did adding more identity categories in the form of microsatellite loci. The addition of an explicit model for the marking process to the categorical SMR also improved the probabilistic identity associations, and was required to estimate density without bias due to the marking process that was not spatially random Whittington *et al.* (2016). One downside of modeling the marking process was that the sparse marking process data introduced more uncertainty into the density estimate of the generalized categorical SMR model, suggesting that generalized SMR estimates could be improved if researchers conduct more substantial marking process surveys, where the expected number of captures for marked individuals is more than that necessary to apply a mark.

We chose our simulation scenarios to be challenging for the conventional SMR model with no telemetry data and no explicit marking process. We are unaware how representative these scenarios are for typical data sets for which SMR is currently being applied. For studies using natural marks, the percentage of marked individuals can be small Jiménez *et al.* (e.g., 2017, individually identified about 10% of the population, but had 2.8 captures/estimated N). Further, for low-density populations with low detectability, recording 0.8 – 1.6 captures per individual as we assumed in the simulations, with this average including those individuals not captured, may be challenging. While we are unaware of any published SMR papers with density estimates that may be biased due to data sparsity and/or a low percentage of marked individuals, our results may explain imprecise and biased density estimates for studies that were not published because they produced implausible or uselessly imprecise density estimates. Further, our simulation trapping array with 81 traps is larger than many remote camera studies, and the effect of trap number, spacing, and extent on unmarked SCR and SMR estimates should be investigated. Preliminary analysis suggested that bias and imprecision are more extreme on trapping arrays with fewer traps and a smaller spatial extent. Given all the factors that may determine the reliability and precision of SMR density estimates, we recommend simulation studies be done before implementing a regular or categorical SMR survey to ensure that the study design is sufficient to produce data that allows for reliable inference.

A second challenge for latent identity SCR models is the presence of individual heterogeneity in detection function parameters, especially *σ*. In both the application of the categorical SPIM (Augustine *et al.*, 2018) and the application of the categorical (conventional) SMR model in this paper, one male individual had an especially large distribution of spatial recaptures, and in both analyses, this one individual’s captures were erroneously split across two individuals when few categorical covariates were used, causing the *σ* estimate to be substantially too low and the density estimate too large. However, our fisher application deviated from typical SMR studies due to the fact that we could not deterministically link marked individual samples during the sighting process (due to the shadow effect; Mills *et al.*, 2000), treating all latent identity samples as unknown marked status samples. If marked individuals could have been identified during sighting, this individual’s samples would have been deterministically linked. In general, as the proportion of marked and identifiable individuals increases, the positive bias due to individual heterogeneity in latent identity SCR models will likely shift to negative bias as is seen in typical SCR models with perfect individual identification.

Given that the typical SCR estimates with perfect individual identification will be conservative (biased low) in the presence of individual heterogeneity in detection function parameters, it is desirable to be able to reproduce these estimates in latent identity models. The categorical SMR model offers more tools over the categorical SPIM to reproduce the null SCR density estimates in the presence of individual heterogeneity in *σ*. When telemetry data or an explicit model for the marking process were included in the fisher analysis, the samples for the individual with a large *σ* were less likely to be split apart, and when they were, the *σ* estimate and density estimates were less affected due to the information coming from these other data sources. Individual heterogeneity could be modeled in latent identity SCR models; however, there is no current consensus about how to do so when it is unknown to what degree compensation between parameters exists (Efford & Mowat, 2014) for any given species and study design. The identity covariate-specific detection functions introduced in this paper offer a solution for the subset of individual heterogeneity in detection function parameters that can be explained by categorical covariates. Finally, when planning latent identity SCR studies, testing the robustness of density estimates to some unmodeled deviations from homogeneity in detection function parameters will help determine the reliability of the density estimates the survey will produce.

While we considered genetic microsatellites in this application, categorical marks are available in other forms of noninvasive capture-recapture methods. As discussed by Augustine *et al.* (2018), when using remote cameras, features such as sex, age class, color morph, natural markings, and/or morphological features may provide categorical identity information. Currently, this information is being used to assign unique individual identities (e.g., Molina *et al.*, 2017; Sirén *et al.*, 2016; Royle *et al.*, 2011); however, not all individuals may have features that permit a unique identification. In this case, ignoring the photographs of these individuals will introduce strong individual heterogeneity in *p*_0_ and thus negative bias in abundance and density estimates. Spatial mark-resight is currently being used to avoid this problem; however, multiple observers can disagree about individual identities (e.g. Rich *et al.*, 2014). The categorical SMR model allows for the photos about which observers agree to be considered as marked with known identities and for the observable features in photos for which some observers thought contained identifiable individuals (e.g., presence of scars, botflies, etc.) to be used as categorical identity covariates, information that is currently being discarded for species including various mustelids (Royle *et al.*, 2011; Sirén *et al.*, 2016), Andean bears (Molina *et al.*, 2017), and pumas (Rich *et al.*, 2014). When applying SMR to naturally marked populations, the number of marked individuals is considered unknown (Royle *et al.*, 2013), which we did not consider in the model development; however, we provide a categorical SMR MCMC sampler for naturally marked populations in Supplement 1.

Similar to Augustine *et al.* (2018), we considered density estimation for populations where all individuals are potentially categorically marked (e.g., sex, age class, and microsatellite loci). The addition of marked individuals in this paper is the first step towards allowing for researcher-deployed categorical marks that only apply to the marked subset of individuals, such as non-unique color tags. If categorical marks are applied to a subset of individuals and the full categorical identities are not unique, their partial identities can be modeled similarly to how the natural partial marks are modeled, except that they only apply to the marked portion of the population. If the numbers of each mark type deployed are known, this information can be used to reduce the uncertainty in individual identity; however constraints on updating the latent identity covariate values will be introduced, requiring a different algorithm to update these values. Researcher-deployed categorical marks would allow for better density estimation for species that can be categorically marked, but the number of each mark types are not sufficient to guarantee each individual has a unique mark. One potential application of categorical marks is the application of colored visual implant elastomers in amphibians. For example, Sutherland *et al.* (2016) marked salamanders in 4 unique locations using 4 colors for a total number of 624 combinations, which might not be perfectly observable upon recapture. Further, if more individuals were marked than the number of unique full categorical identities, some duplicate marks could be deployed and the resulting uncertainty could be incorporated into the density estimate by the categorical SMR model. If all or a subset of marks are applied only to the captured individuals, estimation will be improved over the categorical SPIM and categorical SMR models that consider that uncaptured individuals may have the same full categorical identity as captured individuals.

Finally, we developed efficient MCMC algorithms for conventional and generalized SMR, with and without categorical identity covariates and telemetry data, that allow for all three types of SMR latent identity samples and interspersed marking and sighting occasions. This was done using the 2-dimensional individual by trap data, instead of the 3-dimensional individual by trap by occasion data used to date. The temporal constraints on the allocation of latent identity samples to individuals due to the occasion they were marked as well as the constraints posed by the Bernoulli observation model (can only observe a 0 or 1) are handled using constraint matrices in the latent data update. The additional computational cost of checking the constraint matrices during the latent data update is usually more than offset by the fact that the observation model likelihood, which must be calculated several times per iteration, uses the 2 dimensional data. These MCMC algorithms, as well as a version of conventional SMR for natural marks, where the number of marked individuals is unknown, can be found in Supplement 1. These functions will be maintained in the SPIM package (https://github.com/benaug/SPIM).

## Acknowledgments

Acknowledgments for the fisher data go to: InnoTech Alberta, The Beaver Hills Initiative, NSERC (Canada), MITACS, and Alberta Environment and Parks as major funders. Major acknowledgments go to: M. Pybus, D. Vujnovic, G. Hood, B. Eaton, M. McAdie, I. Brusselers, T. Zembal and Wildlife Genetics International for aid in research facilitation, data collection, and genetic analyses.

## 7 Appendix A: MCMC algorithm

We begin by restating some of the notation of the categorical SMR model. As in the Methods section, we will describe a generalized SMR version with a marking and a resighting process. The conventional SMR model version is identical to the generalized SMR version except there is no marking process. We assume individuals are marked at locations ***X**^M^* (dimension *J^M^* × 2) over *K^M^* occasions and resighted at locations ***X**^S^* (dimension *J^S^* × 2) over *K^S^* occasions. The observed marking process capture history is ***Y**^M^* of dimension *n^M^* × *J^M^*, where *n^M^* is the number of marked individuals, a subset of the unknown total number of individuals captured and/or sighted, *n^cap^.* The *n^cap^* × *J^S^* sighting history, ***Y**^S^*, is divided across up to four observed data matrices: ***Y**^S.M^*, the sighting history of the marked and identified individuals, ***Y**^S.M.unk^*, the sighting history of the marked and unidentified individuals, ***Y**^S.um^*, the sighting history of the unmarked individuals, and ***Y**^S.unk^*, the sighting history for the unknown marked status individuals. These latter three matrices contain the latent identity samples with *J^S^* columns and *n^M.unk^*, *n^um^*, and *n^unk^* rows, respectively. Each row of these matrices, corresponding to a single observation, consists of all zero entries except for a single “1” indicating the trap of capture. For illustration, we reproduce the hypothetical true and observed data matrices from the Methods for two unmarked individuals captured at *J* = 3 traps. The first individual was captured once in the first trap and once in the thrid trap, while the second individual was captured twice in the first trap and once each in the second and third traps. In this example, the marked status (unmarked) was observed for all but the 2^nd^ individual’s captures in traps 2 and 3. Each single observation in ***Y**^S^* is then disaggregated into the rows of the appropriate latent identity observation matrix, ***Y**^S.um^* or ***Y**^S.unk^*, depending on whether the marked status is observed for these two unmarked individuals.

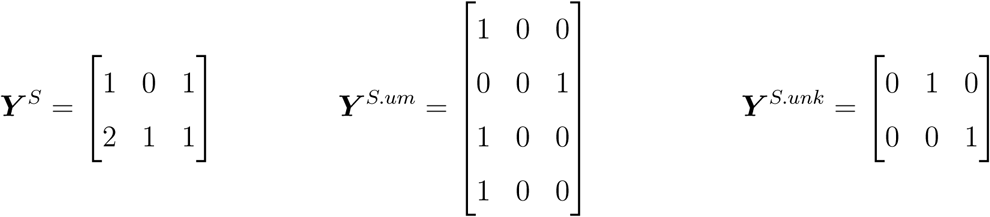

The marked individuals’ latent identity samples are similarly disaggregated into ***Y**^S.M.unk^* or ***Y**^S.unk^*, also depending on whether their marked status can be determined. The ordering of the latent identity samples does not matter – the only information being used to indicate their identity is the trap of capture linked with their observed categorical identity.

Next, we define the *N* × *n^cat^* matrix ***G*** to be the categorical identity matrix whose rows correspond to the rows of ***Y**^S^*, with each column storing the values of each categorical covariate. The observed categorical identities are stored in ***G**^M^*, ***G**^M.unk^*, ***G**^um^*, and ***G**^unk^*, with the number of rows matching their corresponding latent identity sighting histories, each with *n^cat^* columns. The categorical identities for the marked individuals are recorded in ***G**^M^*, whose rows correspond to those in the marked individual sighting history, ***Y**^S.M^*. The categorical identities for the latent identity samples are stored in ***G**^M.unk^*, ***G**^um^*, or ***G**^unk^*, depending on sample type, whose rows correspond to their appropriate latent identity sighting history, ***Y**^S.M.unk^*, ***Y**^S.um^* or ***Y**^S.unk^*. The example categorical identity matrices presented in the Methods section corresponding to the true and observed capture histories for two unmarked individuals above are:

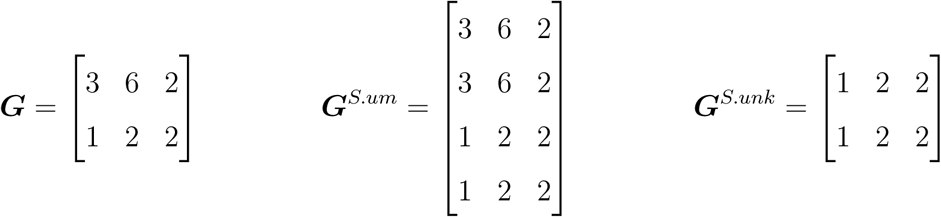

In this scenario, individual 1 has full categorical identity “3, 6, 2” and individual 2 has full categorical identity “1, 2, 2”. Because the marked status for both of the first individual’s samples was observed, its observed categorical identity is recorded in the first two rows of ***G**^S.um^*, corresponding to the first two rows of ***G**^S.um^*. The marked status of the second individual’s captures in trap 1 was observed, so its observed categorical identity is recorded in the third and fourth rows of ***G**^S.um^*, corresponding to the second two rows of ***G**^S.um^*. Then, the marked status of the second individual’s captures in traps 2 and 3 was not observed, so its observed categorical identity is recorded in the first two rows of ***G**^S.unk^*, corresponding to the first two rows of ***Y**^S.unk^*. The categorical covariates for the latent identity samples for the marked individuals are similarly stored in *G^S.M.unk^* or ***G**^S.unk^*, depending on whether their marked status is observed. In this example, all identity covariate values were observed, but missing observations are allowed, coded as “0”.

The partially latent, true sighting history and categorical identity matrix, ***Y**^S.true^* and ***G**^true^*, are initialized by augmenting ***Y**^S.M^* and ***G**^M^* up to dimension *M* × *J^S^* and *M* × *n^cat^*, respectively, with all zero entries, where *M* is the level of data augmentation. The marking process capture history, ***Y**^M^*, is also augmented up to dimension *M* × *J^M^*. The latent elements of ***Y**^S.true^* are then initialized by allocating to it the samples in the latent identity observed capture histories using spatial proximity and consistency between their observed categorical identities. The complete or minimally-implied categorical identity for each row of ***Y**^S.true^* is stored in the corresponding row of ***G**^true^*. If each identity covariate is observed for at least one set of linked samples, the full categorical identity will be complete, while the categorical identity will be partially latent if the value of one or more covariates is not observed for any of the liked samples. Finally, the latent elements of ***G**^true^*, whether the result of missing identity covariate data or corresponding to the augmented individuals, are initialized using the starting values for the population category level frequencies following 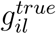 ∼ Categorical(*γ*_1_) for each of the *l* categorical identity covariates. The inferential process is to probabilistically reconstruct the latent elements of ***Y**^S.true^* and ***G**^true^*, estimate the activity centers, ***S***, the elements of the data augmentation indicator vector ***z***, the data augmentation inclusion probability *ϕ*, the population category level frequencies *γ*, the category level-specific detection function parameters for both observation processes, 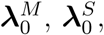 and ***σ***, and then abundance and density.

In a mark-resight survey where the marking and sighting occasions are interspersed, the marked status of each individual may change across sighting occasions (for example, an individual is unmarked at occasion 1 and then becomes marked on a subsequent occasion). In order to allow for the marking and sighting processes to overlap while using the 2-dimensional individual by trap level data that excludes the occasion dimension, we use the 3-dimensional individual by trap by occasion data to construct indicator matrices. These constraint matrices either allow or disallow latent identity samples to be allocated to each marked individual depending on the occasion that each individual was marked and the occasion on which the latent identity samples were observed. The constraint matrices are ***C**^M.unk^* and ***C**^um^*, with *n^M^* rows and *n^M.unk^* and *n^um^* columns, respectively. No constraint matrix for unknown marked status samples is required because they can be allocated to individuals with or without a mark on any occasion. Further, the constraint matrices do not need to account for the individuals that are never marked because their marked status never changes. Each constraint matrix has elements “1” if latent sample *j* can be combined with marked individual *i* and elements “0” otherwise.

These constraint matrices can be built using the observed occasions of the latent identity observations and following Whittington *et al.* (2016), a *n^M^* × *K* indicator matrix, ***m***, indicating the occasions that each individual was marked with a “1” and “0” otherwise. For example, consider three individuals where individual one was marked (marked status of marked) on all three occasions, individual two was marked on the second two occasions, and individual three was marked on the third occasion. Then, consider two unknown marked status samples collected on occasions 1 and 2 and two marked status but unknown identity samples collected on occasions 1 and 2. The resulting ***m*** and constraint matrices would be:

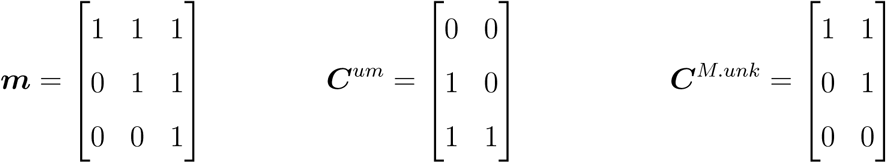

The first unmarked sample cannot belong to the first individual, who is marked on the first occasion when the sample was collected. Similarly, the second unmarked sample cannot belong to the first two individuals, who were marked on the second occasion when the sample was collected. Conversely, the first marked sample cannot belong to individuals two and three and the second marked sample cannot belong to the third individual. If the sighting process is binomial, another constraint matrix is used to disallow latent captures to be combined in a manner that produce individual by trap by occasion observations greater than 1 (described in Augustine *et al.*, 2018). Finally, we omit the description of telemetry data here, referring readers to Whittington *et al.* (2016), which describes how we accommodated telemetry data in the categorical SMR model.

The joint posterior is then:

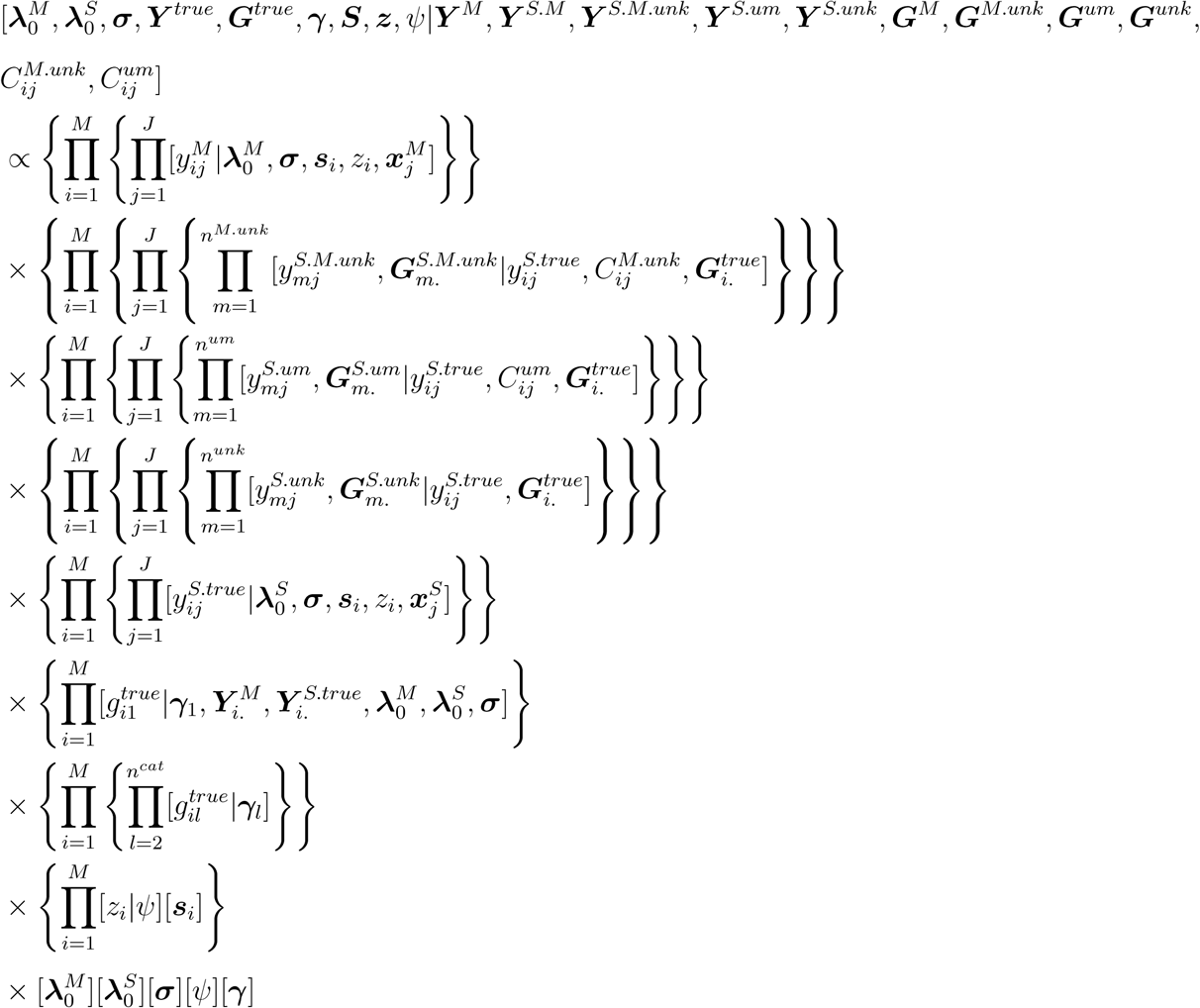

We assume the following prior distributions:

1. 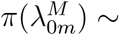 Uniform(0, ∞), for 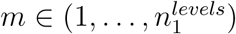
2. 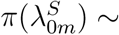Uniform(0, ∞), for 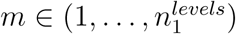
3. 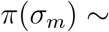Uniform(0, ∞), for 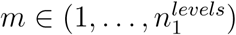
4. 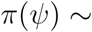Uniform(0,1)
5. 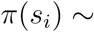Uniform(𝓢)
6. *π*(*γ_l_*) ∼ Dirichlet(*α_l_*), where ***α**_l_* is the vector of Dirichlet parameters of length 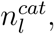 indexed by *g* below. All ***α**_l_* were set to vectors of 1.

Here, we list the full conditional distributions or the distributions they are proportional to. Note, the first six components are exactly the same as a typical SCR model except there are two observation processes, the seventh and eighth component correspond to the categorical identity model, and the final component corresponds to the uncertain identity model. We omit notation indicating that the detection function parameter distributions are conditional on their respective capture locations, *X^**M**^* and ***X**^S^*, which determine the individual by trap encounter rates or probabilities.

1. Baseline encounter rate for the marking process that varies by one categorical identity covariate. 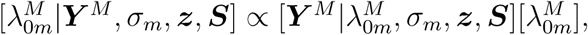 where 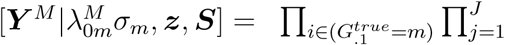 Binomial 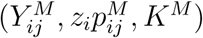 for the Bernoulli observation model and 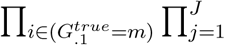 Poisson 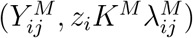 for the Poisson observation model. 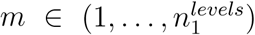
2. Baseline encounter rate for the sighting process that varies by one categorical identity covariate. 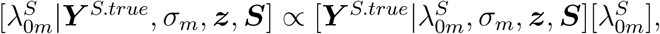 where 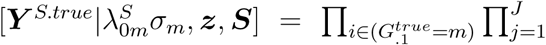 Binomial 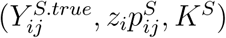 for the Bernoulli observation model and 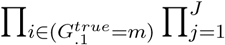 Poisson 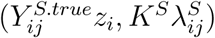 for the Poisson observation model. 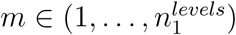
3. The spatial scale parameter that varies by one categorical identity covariate, shared by the marking and sighting processes. 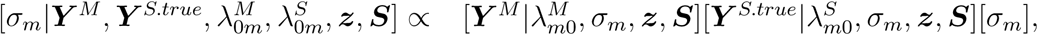 where 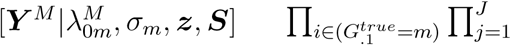 Binomial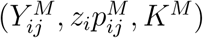 for the Bernoulli observation model and 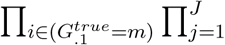 Poisson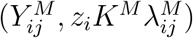 for the Poisson observation model and 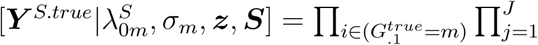 Binomial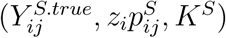 for the Bernoulli observation model. Then, 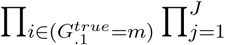 Poisson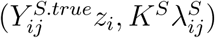 for the Poisson observation model. 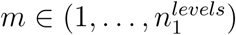
4. The data augmentation indicator vector. 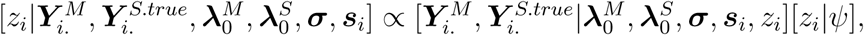 where 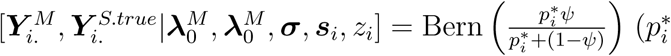 defined below in step C of the MCMC algorithm)
5. The data augmentation inclusion probability. [*ψ*|***z***] = Beta(1 + Σ_*i*_ *z_i_*, 1 + *M* – Σ_*i*_ *z_i_*)
6. The activity centers. 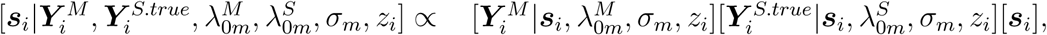 where 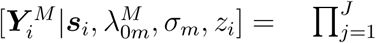 Binomial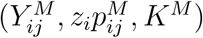 for the Bernoulli observation model and 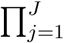 Poisson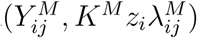 for the Poisson observation model. and 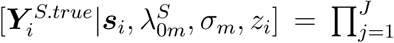 Binomial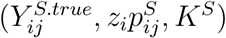 for the Bernoulli observation model and 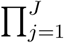 Poisson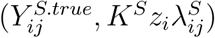 for the Poisson observation model. 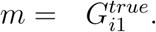
7. The categorical identity covariate upon which the detection function parameters depend (assumed to be the first categorical covariate). 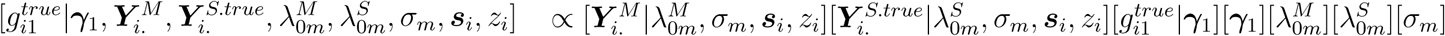 where 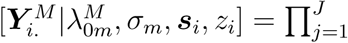 Binomial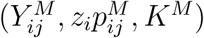 for the Bernoulli observation model and 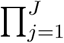 Poisson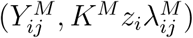 for the Poisson observation model. Then, 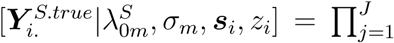 Binomial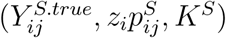 for the Bernoulli observation model and 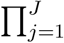 Poisson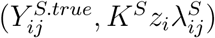 for the Poisson observation model. 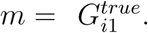 Then, 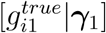 = Categorical(*γ*_1_).
8. The categorical identity covariates that do not determine the detection function parameters. 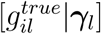 = Categorical(*γ_l_*) for *l* = 2,…, *n^cat^*
9. The categorical identity covariate level probabilities 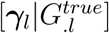 = Dirichlet(***y**_l_* + ***α**_l_*) for all *l*, where *y_l_g__* is the number of individuals in the population (*z_i_* = 1) with categorical identity *g* at loci *l* and ***α**_l_* is the Dirichlet prior for *γ_l_*, a vector of 1’s.
10. The latent data distributions and associated observed categorical identities. The distribution for the *m^th^* marked but unknown identity sample at trap *j* and the resulting observed categorical identity depends on the true sighting history at trap *j*, the identity constraint matrix for the marked but unknown identity samples, and the true full categorical identities. 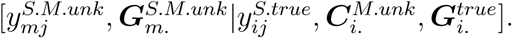 Similarly, 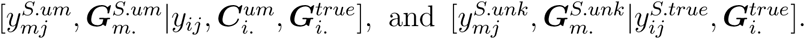 The full conditional distributions for each of these model components is the unmarked SCR multinomial full conditional (Chandler & Royle, 2013) with different cell probabilities zeroed out as described below in MCMC step F.

Here we outline the MCMC algorithm. Again, note that the first five steps are exactly the same as a typical SCR model, but with two observation processes.

A. Update λ_o*m*_ for the marking and sighting processes. λ_o*m*_ is updated with a Metropolis-Hastings step using the distribution Normal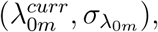 to propose 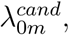 automatically rejecting if a negative value is proposed. The MH ratio is computed using distributions 1 or 2 above, for the marking and sighting processes, respectively.
B. Update *σ_m_*. *σ_m_* is updated with a Metropolis-Hastings step using the distribution Normal(*σ^curr^*,*σ_σ_m__*), to propose 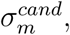 automatically rejecting if a negative value is proposed. The MH ratio is computed using distribution 3 above.
C. Update **z**. Each *z_i_* is updated by a Gibbs step using the full conditional above where 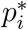 is the probability individual *i* was not captured during the both observation processes. Let *p̄_ij_* be the probability of not being captured for each individual at each trap over the *K^M^* marking and *K^S^* sighting occasions. For the bernoulli observation process, this is 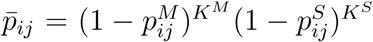 and for the Poisson observation process, we first compute 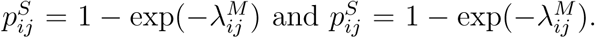 The probability of not being captured during the experiment for each individual is then 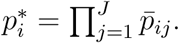
D. Update *Ψ*. *Ψ* is updated with a Gibbs step. Since *π*(*Ψ*) ∼ Uniform(0,1) is in the Beta family, the full conditional distribution for *Ψ* is Beta(1 + Σ_*i*_ *z_i_*, 1 + *M* – Σ_*i*_*z_i_*).
E. Update **s**. Each activity center ***s**_i_* is updated with a Metropolis-Hastings step using the distributions Normal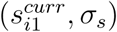 and Normal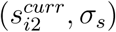 to propose 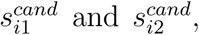 respectively. Proposals that fall outside of the state-space are rejected. The MH ratio is calculated using distribution 6 above.
F. Update ***Y**^S.true^.* To update ***Y**^S.true^* for the Poisson observation model, we use a class-structured version of the algorithm used by Whittington *et al.* (2016), but update the latent identity samples one by one. If there is only one identity class as in typical SMR, the full conditional for the focal latent identity sample *f* observed at trap *j* is:

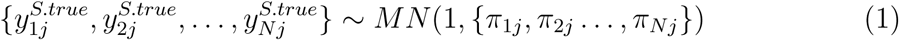

with *π_ij_* = *z_i_*λ_*ij*_/Σ_*i*_ *z_i_*λ_*i*_, and different elements of λ*_ij_* set to zero depending on the observed sample type. For unknown marked status samples, no elements of λ*_ij_* are set to zero. For marked but unknown identity samples, elements *n^M^* + 1,…, *M* are zeroed out. Then, for unmarked samples, elements 1,…, *n^M^* are zeroed out. When the marking and sighting processes overlap, the compatibility of the focal sample must be checked with all of the marked individuals using the constraint matrices specified above. Any of the *i* individuals whose marked status is inconsistent with the *f* focal sample have their λ*_ij_* values set to zero prior to the calculation of *π_ij_*. When categorical identity covariates are introduced, further restrictions are placed on λ*_ij_*. Any individuals whose full categorical identity does not match that of the focal sample have their λ*_ij_* values set to zero prior to the calculation of *π_ij_*. In all cases, once *π_ij_* is calculated with the appropriate elements set to zero, the focal sample, *f* is updated in *Y^S.true^* using the full conditional described above. For the Bernoulli observation model, the multinomial full conditional result above does not hold so we use a Metropolis-Hastings proposal. We use the multinomial full conditional above with the appropriate constraints depending on sample type and full categorical identity as the proposal distribution. Because the transition distribution for selecting samples to update is not symmetric, the differing forward and backwards transition probabilities must be calculated. The forward transition probability is the probability of selecting a sample to update and then the probability of choosing the new identity 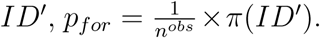 The backwards transition probability accounting for the fact that you can only select individuals with full categorical identities compatible with the focal observed categorical identity to move back is 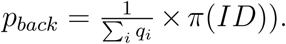 We then accept the proposal with probability:

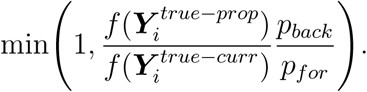

where *f* (.) is the SCR sighting observation model likelihood.
G. Update 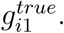 Elements of 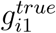 are updated one at a time, using the distribution 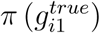 Categorical(*γ_l_*) to propose 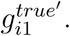 We then accept the proposal with probability

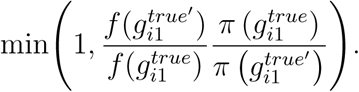

where f(.) is distribution 7 above.
H. Update 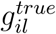 for *l* = 2,…, 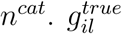 is updated for all elements that are not determined by the samples currently allocated to individual *i* using distribution 8 listed above.
I. Update *γ_l_*. *γ_l_* is updated with a Gibbs step. Following Wright et al. (2009), we adopt a Dirichlet prior for *γ_l_*, leading to a Dirichlet full conditional with parameter vector 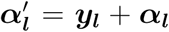 where *y_lg_* is the number of individuals in the population with category level *g* for covariate *l*. To draw values from the full conditional, we first simulate a vector of Gamma random variables ***g**_l_* ∼ Gamma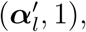 where ***g**_l_* is of length 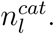 Then, renormalizing these gamma random variables is a draw from the Dirichlet full conditional, i.e. 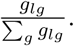
J. Calculate the derived quantities population abundance, 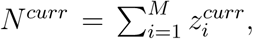 population density, 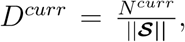 and the number of unique individuals captured 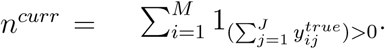 The number of unique marked and unmarked individuals captured may also be computed.

## Appendix B: 8 Simulation A Results

**Table 3:**
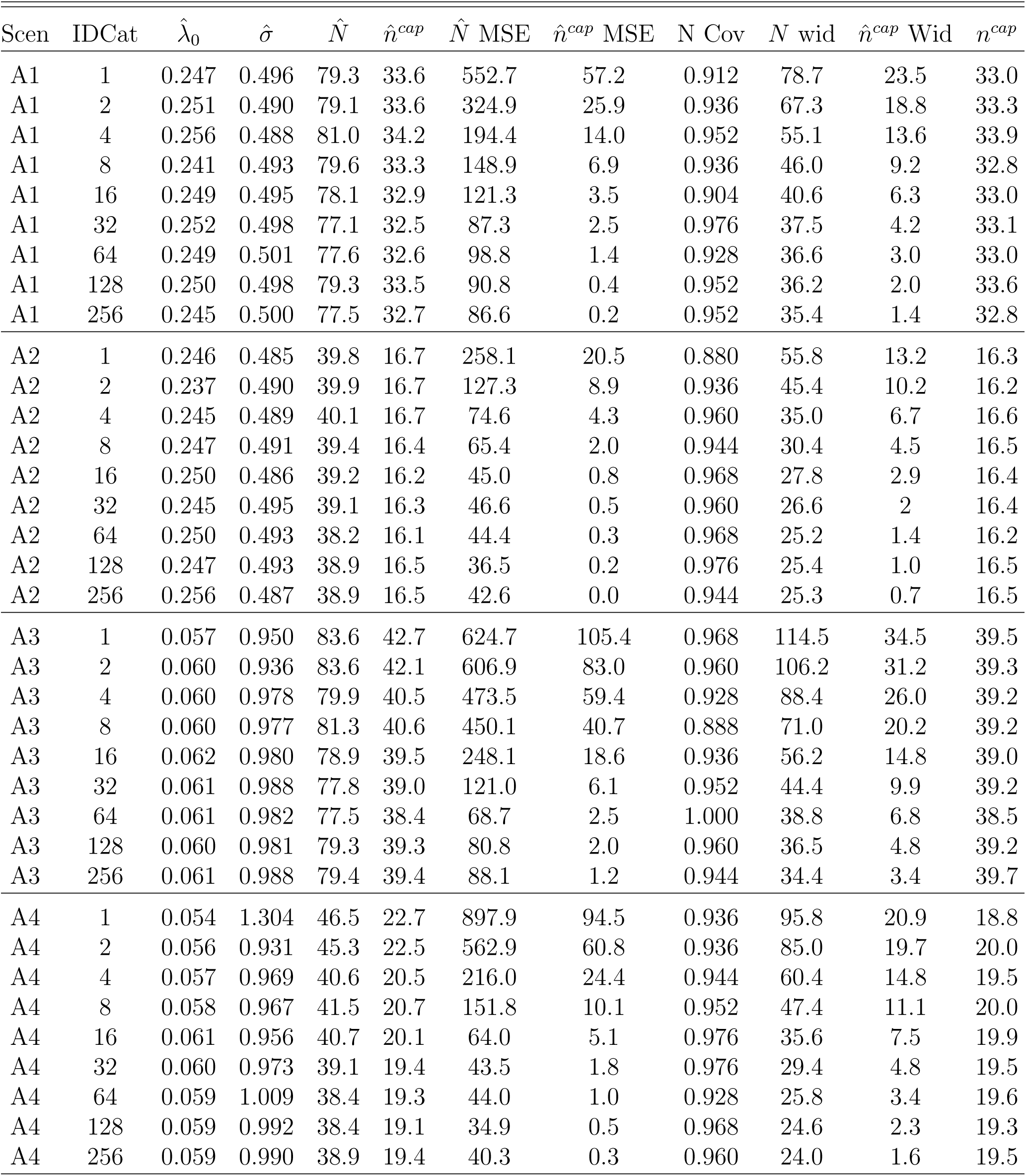
Results from Simulation A Scenarios 1-4 where 10% of the population was marked. The columns are “IDCat”, the number of identity categories, the mean point estimates, the 95% CI coverage of abundance, the mean width of the 95% CIs for N and *n^um^*, and the mean number of simulated individuals that were captured.

| Scen | IDCat | $\hat{\lambda}_0$ | $\hat{\sigma}$ | $\hat{N}$ | $\hat{n}^{cap}$ | $\hat{N}$ MSE | $\hat{n}^{cap}$ MSE | N Cov | $N$ wid | $\hat{n}^{cap}$ Wid | $n^{cap}$ |
| --- | --- | --- | --- | --- | --- | --- | --- | --- | --- | --- | --- |
| A1 | 1 | 0.247 | 0.496 | 79.3 | 33.6 | 552.7 | 57.2 | 0.912 | 78.7 | 23.5 | 33.0 |
| A1 | 2 | 0.251 | 0.490 | 79.1 | 33.6 | 324.9 | 25.9 | 0.936 | 67.3 | 18.8 | 33.3 |
| A1 | 4 | 0.256 | 0.488 | 81.0 | 34.2 | 194.4 | 14.0 | 0.952 | 55.1 | 13.6 | 33.9 |
| A1 | 8 | 0.241 | 0.493 | 79.6 | 33.3 | 148.9 | 6.9 | 0.936 | 46.0 | 9.2 | 32.8 |
| A1 | 16 | 0.249 | 0.495 | 78.1 | 32.9 | 121.3 | 3.5 | 0.904 | 40.6 | 6.3 | 33.0 |
| A1 | 32 | 0.252 | 0.498 | 77.1 | 32.5 | 87.3 | 2.5 | 0.976 | 37.5 | 4.2 | 33.1 |
| A1 | 64 | 0.249 | 0.501 | 77.6 | 32.6 | 98.8 | 1.4 | 0.928 | 36.6 | 3.0 | 33.0 |
| A1 | 128 | 0.250 | 0.498 | 79.3 | 33.5 | 90.8 | 0.4 | 0.952 | 36.2 | 2.0 | 33.6 |
| A1 | 256 | 0.245 | 0.500 | 77.5 | 32.7 | 86.6 | 0.2 | 0.952 | 35.4 | 1.4 | 32.8 |
| A2 | 1 | 0.246 | 0.485 | 39.8 | 16.7 | 258.1 | 20.5 | 0.880 | 55.8 | 13.2 | 16.3 |
| A2 | 2 | 0.237 | 0.490 | 39.9 | 16.7 | 127.3 | 8.9 | 0.936 | 45.4 | 10.2 | 16.2 |
| A2 | 4 | 0.245 | 0.489 | 40.1 | 16.7 | 74.6 | 4.3 | 0.960 | 35.0 | 6.7 | 16.6 |
| A2 | 8 | 0.247 | 0.491 | 39.4 | 16.4 | 65.4 | 2.0 | 0.944 | 30.4 | 4.5 | 16.5 |
| A2 | 16 | 0.250 | 0.486 | 39.2 | 16.2 | 45.0 | 0.8 | 0.968 | 27.8 | 2.9 | 16.4 |
| A2 | 32 | 0.245 | 0.495 | 39.1 | 16.3 | 46.6 | 0.5 | 0.960 | 26.6 | 2 | 16.4 |
| A2 | 64 | 0.250 | 0.493 | 38.2 | 16.1 | 44.4 | 0.3 | 0.968 | 25.2 | 1.4 | 16.2 |
| A2 | 128 | 0.247 | 0.493 | 38.9 | 16.5 | 36.5 | 0.2 | 0.976 | 25.4 | 1.0 | 16.5 |
| A2 | 256 | 0.256 | 0.487 | 38.9 | 16.5 | 42.6 | 0.0 | 0.944 | 25.3 | 0.7 | 16.5 |
| A3 | 1 | 0.057 | 0.950 | 83.6 | 42.7 | 624.7 | 105.4 | 0.968 | 114.5 | 34.5 | 39.5 |
| A3 | 2 | 0.060 | 0.936 | 83.6 | 42.1 | 606.9 | 83.0 | 0.960 | 106.2 | 31.2 | 39.3 |
| A3 | 4 | 0.060 | 0.978 | 79.9 | 40.5 | 473.5 | 59.4 | 0.928 | 88.4 | 26.0 | 39.2 |
| A3 | 8 | 0.060 | 0.977 | 81.3 | 40.6 | 450.1 | 40.7 | 0.888 | 71.0 | 20.2 | 39.2 |
| A3 | 16 | 0.062 | 0.980 | 78.9 | 39.5 | 248.1 | 18.6 | 0.936 | 56.2 | 14.8 | 39.0 |
| A3 | 32 | 0.061 | 0.988 | 77.8 | 39.0 | 121.0 | 6.1 | 0.952 | 44.4 | 9.9 | 39.2 |
| A3 | 64 | 0.061 | 0.982 | 77.5 | 38.4 | 68.7 | 2.5 | 1.000 | 38.8 | 6.8 | 38.5 |
| A3 | 128 | 0.060 | 0.981 | 79.3 | 39.3 | 80.8 | 2.0 | 0.960 | 36.5 | 4.8 | 39.2 |
| A3 | 256 | 0.061 | 0.988 | 79.4 | 39.4 | 88.1 | 1.2 | 0.944 | 34.4 | 3.4 | 39.7 |
| A4 | 1 | 0.054 | 1.304 | 46.5 | 22.7 | 897.9 | 94.5 | 0.936 | 95.8 | 20.9 | 18.8 |
| A4 | 2 | 0.056 | 0.931 | 45.3 | 22.5 | 562.9 | 60.8 | 0.936 | 85.0 | 19.7 | 20.0 |
| A4 | 4 | 0.057 | 0.969 | 40.6 | 20.5 | 216.0 | 24.4 | 0.944 | 60.4 | 14.8 | 19.5 |
| A4 | 8 | 0.058 | 0.967 | 41.5 | 20.7 | 151.8 | 10.1 | 0.952 | 47.4 | 11.1 | 20.0 |
| A4 | 16 | 0.061 | 0.956 | 40.7 | 20.1 | 64.0 | 5.1 | 0.976 | 35.6 | 7.5 | 19.9 |
| A4 | 32 | 0.060 | 0.973 | 39.1 | 19.4 | 43.5 | 1.8 | 0.976 | 29.4 | 4.8 | 19.5 |
| A4 | 64 | 0.059 | 1.009 | 38.4 | 19.3 | 44.0 | 1.0 | 0.928 | 25.8 | 3.4 | 19.6 |
| A4 | 128 | 0.059 | 0.992 | 38.4 | 19.1 | 34.9 | 0.5 | 0.968 | 24.6 | 2.3 | 19.3 |
| A4 | 256 | 0.059 | 0.990 | 38.9 | 19.4 | 40.3 | 0.3 | 0.960 | 24.0 | 1.6 | 19.5 |

**Table 4.**

| Scen | IDCat | $\hat{\lambda}_0$ | $\hat{\sigma}$ | $\hat{N}$ | $\hat{n}^{cap}$ | $\hat{N}$ MSE | $\hat{n}^{cap}$ MSE | N Cov | $N$ wid | $\hat{n}^{cap}$ Wid | $n^{cap}$ |
| --- | --- | --- | --- | --- | --- | --- | --- | --- | --- | --- | --- |
| A1c | 1 | 0.120 | 0.471 | 83.4 | 25.1 | 1,074.7 | 30.4 | 0.936 | 102.1 | 15.2 | 23.2 |
| A1c | 2 | 0.124 | 0.465 | 84.3 | 25.2 | 662.5 | 14.2 | 0.960 | 87.7 | 12.6 | 24.0 |
| A1c | 4 | 0.120 | 0.483 | 81.9 | 25.0 | 310.7 | 8.7 | 0.968 | 74.0 | 10.0 | 24.2 |
| A1c | 8 | 0.124 | 0.482 | 80.2 | 24.4 | 259.7 | 4.2 | 0.944 | 59.9 | 7.2 | 24.3 |
| A1c | 16 | 0.123 | 0.487 | 78.5 | 23.7 | 200.8 | 2.2 | 0.952 | 52.6 | 5.1 | 23.8 |
| A1c | 32 | 0.122 | 0.487 | 78.7 | 23.9 | 107.0 | 1.1 | 0.992 | 49.3 | 3.6 | 24.1 |
| A1c | 64 | 0.125 | 0.491 | 78.1 | 23.9 | 116.6 | 0.6 | 0.968 | 45.9 | 2.4 | 24.1 |
| A1c | 128 | 0.123 | 0.485 | 78.2 | 23.7 | 122.9 | 0.4 | 0.976 | 45.4 | 1.6 | 23.9 |
| A1c | 256 | 0.130 | 0.485 | 79.1 | 24.3 | 124.5 | 0.3 | 0.936 | 44 | 1.2 | 24.4 |
| A2c | 1 | 0.105 | 0.471 | 47.0 | 13.6 | 653.7 | 12.7 | 0.944 | 90.2 | 9.1 | 12.4 |
| A2c | 2 | 0.114 | 0.453 | 44.6 | 13.0 | 342.1 | 6.5 | 0.960 | 77.6 | 7.2 | 12.3 |
| A2c | 4 | 0.121 | 0.468 | 39.6 | 12.0 | 90.4 | 2.1 | 0.968 | 49.9 | 5.0 | 11.9 |
| A2c | 8 | 0.123 | 0.487 | 39.1 | 12.0 | 66.8 | 1.1 | 0.968 | 40.6 | 3.6 | 12.0 |
| A2c | 16 | 0.122 | 0.479 | 40.1 | 12.0 | 85.5 | 0.6 | 0.944 | 38.6 | 2.4 | 12.2 |
| A2c | 32 | 0.123 | 0.483 | 40.1 | 12.2 | 65.5 | 0.4 | 0.960 | 35.6 | 1.6 | 12.4 |
| A2c | 64 | 0.123 | 0.479 | 40.0 | 12.2 | 75.1 | 0.3 | 0.952 | 35.2 | 1.2 | 12.3 |
| A2c | 128 | 0.127 | 0.475 | 38.9 | 11.8 | 56.5 | 0.1 | 0.952 | 33.1 | 0.8 | 11.9 |
| A2c | 256 | 0.123 | 0.484 | 38.6 | 11.8 | 61.3 | 0.1 | 0.928 | 32.4 | 0.4 | 11.9 |
| A3c | 1 | 0.028 | 0.959 | 84.6 | 27.4 | 1,304.2 | 34.0 | 0.928 | 113.8 | 16.7 | 25.7 |
| A3c | 2 | 0.027 | 0.938 | 82.7 | 27.5 | 555.1 | 23.7 | 0.968 | 111.8 | 15.6 | 25.6 |
| A3c | 4 | 0.029 | 0.917 | 84.1 | 27.8 | 686.4 | 23.2 | 0.968 | 103.3 | 14.2 | 26.5 |
| A3c | 8 | 0.031 | 0.928 | 81.7 | 27.2 | 523.2 | 14.7 | 0.960 | 85.0 | 11.8 | 26.6 |
| A3c | 16 | 0.030 | 0.918 | 84.1 | 27.6 | 391.9 | 9.1 | 0.952 | 77.6 | 9.2 | 26.3 |
| A3c | 32 | 0.031 | 0.910 | 81.8 | 26.4 | 230.5 | 3.6 | 0.968 | 66.1 | 6.7 | 26.1 |
| A3c | 64 | 0.031 | 0.928 | 79.8 | 26.1 | 214.1 | 1.8 | 0.968 | 56.0 | 4.8 | 26.0 |
| A3c | 128 | 0.030 | 0.961 | 77.5 | 25.7 | 143.0 | 1.2 | 0.976 | 51.2 | 3.4 | 25.8 |
| A3c | 256 | 0.029 | 0.989 | 77.8 | 26.0 | 141.2 | 0.6 | 0.944 | 48.8 | 2.4 | 26.2 |
| A4c | 1 | 0.028 | 0.883 | 46.0 | 14.9 | 716.9 | 14.0 | 0.952 | 99.2 | 10.8 | 13.7 |
| A4c | 2 | 0.028 | 1.883 | 45.8 | 14.8 | 606.4 | 14.9 | 0.928 | 96.5 | 9.5 | 13.3 |
| A4c | 4 | 0.030 | 0.908 | 45.8 | 14.6 | 604.0 | 9.3 | 0.936 | 86.4 | 8.2 | 13.5 |
| A4c | 8 | 0.026 | 0.904 | 45.2 | 14.1 | 376.4 | 6.8 | 0.952 | 78.4 | 6.8 | 12.9 |
| A4c | 16 | 0.029 | 0.912 | 41.4 | 13.6 | 107.9 | 2.6 | 0.968 | 59.6 | 5.2 | 13.2 |
| A4c | 32 | 0.029 | 1.369 | 40.3 | 13.2 | 107.0 | 1.6 | 0.960 | 49.4 | 3.5 | 13.2 |
| A4c | 64 | 0.030 | 0.925 | 40.1 | 13.1 | 76.5 | 0.8 | 0.976 | 42.5 | 2.5 | 13.2 |
| A4c | 128 | 0.030 | 0.964 | 38.8 | 12.9 | 76.0 | 0.4 | 0.968 | 38.7 | 1.6 | 12.9 |
| A4c | 256 | 0.028 | 0.950 | 38.2 | 12.7 | 90.2 | 0.2 | 0.936 | 36.5 | 1.2 | 12.7 |

